# The infection cycle of the haloarchaeal virus HFTV1 is tightly regulated and strongly inhibits motility of its host

**DOI:** 10.1101/2025.04.24.650469

**Authors:** Sabine Schwarzer, Leonard E Bäcker, Jeroen G Nijland, Ismail Hayani Aji, Anne de Jong, Cristina Moraru, Claudia Steglich, Tessa EF Quax

## Abstract

Although viruses have been shown to infect all domains of life, our understanding of the genetic program behind the exploitation of host resources to produce progeny virions is thus far limited to several bacterial viruses. Therefore, to elucidate the transcriptome of euryarchaeal viruses and their hosts, we employed RNAseq analysis of samples taken at different time points from *Haloferax gibbonsii* LR2-5 cultures infected with the lytic model virus Haloferax Tailed Virus 1 (HFTV1). While following the transcription of viral genes throughout the infective life cycle, we observed a tight temporal regulation of viral transcripts as well as differential expression from within viral gene clusters. Furthermore, anti-sense RNAs (asRNAs) appear to play an important role in support of the timing of late-expressed viral genes. Therefore, with many differentially expressed transcripts, including intragenic transcripts and asRNAs, the regulatory machinery employed by HFTV1 contrasts with viral model systems (based on phages), in which antitermination and/or alternative polymerases (seemingly lacking in HFTV1) are more widespread. When looking into differentially expressed host genes, we observed a strong downregulation of genes involved in motility, such as the archaellum and chemotaxis machinery, which was confirmed with swimming assays of HFTV1 infected cells. This might be a strategy of the virus to redirect energy flowing into movement towards the production of virions. In conclusion, this work thus provides a stepping stone for further exploration of the intriguing strategies of viral transcriptional regulation of their infection cycle across the domains of life.

**IMPORTANCE:** Viruses infect members of all three domains of life, including *Archaea*. *Euryarchaea* are widespread microorganisms found in various environments such as the human gut and solar salterns. Due to the exceptional availability of cell biology and genetic tools of some salt-loving archaea, they are a model system to extrapolate from. Insights into the regulation of viral infections are of particular importance, especially since HFTV1, has been adopted as a model virus by the archaeal viral community.

We found that, while harboring parallels with bacterial viruses, such as tight temporal regulation, HFTV1 harbors an impressive number of differentially expressed transcriptional units. Furthermore, anti-sense RNAs and intragenic regulatory elements seem to play a much more prominent role in HFTV1 gene expression. Thus, this work challenges current models and provides valuable new insights into the gene regulation of viral infection of archaea, which mark similarities and differences with viruses from other domains of life.

## INTRODUCTION

Viral predation has been a continuous contributor to evolution and has brought forward countless host adaptions. Archaea, which are a diverse group of microorganisms that are more related with eukaryotes than with bacteria, have likewise been found to be preyed upon by viruses. These viruses are characterized by a high diversity of their genome content and viral capsids (Prangishvili et al., 2017). The study of archaeal viral diversity and infection mechanisms has been important to draw hypotheses about the origin and evolutionary trajectory of viruses in general (Prangishvili et al., 2017). Research on archaeal viruses lags behind our understanding of eukaryotic and bacterial viruses, due to the extreme conditions in which some hosts thrive and the lack of well-developed genetically accessible model virus-host systems for archaea. For example, details of transcriptional regulation of infection are not available for most archaeal viruses. To bridge this knowledge gap, we selected

*Haloferax gibbonsii* LR2-5 and its virus Haloferax tailed virus 1 (HFTV1), as these are a model for the study of virus-host interactions in *Euryarchaea* (Tittes et al. 2021). *H. gibbonsii* LR2-5 and HFTV1 were isolated together from the hypersaline Lake Retba in Senegal (Mizuno et al., 2019). Besides HFTV1, nine other haloarchaeal viruses were found to infect *H. gibbonsii* LR2-5, making it a versatile haloarchaeal viral host (Aguirre Sourrouille et al., 2022). *H. gibbonsii* LR2-5 is closely related to *H. volcanii*. Whereas the *H. volcanii* genome contains several anti-viral defense systems, such as CRISPR-cas, the *H. gibbonsii* genome is almost completely devoid of known defence systems (Tittes et al., 2021). This could be a possible reason for its high viral susceptibility in comparison to *H. volcanii* (Aguirre Sourrouille et al., 2022).

*H. gibbonsii* LR2-5 displays motility behavior in liquid medium and on semi-solid agar plates (Tittes et al., 2021). Cells swim forward and reverse with the help of the archaellum, the rotating motility structure of archaea. By adjusting their forward and backwards runs they can swim towards or away from environmental stimuli, which are sensed by the chemotaxis system. In addition, *H. gibbonsii* LR2-5 was also shown to display growth-phase dependent changes in cell shape, as has been reported for multiple haloarchaea (Schwarzer et al., 2021; Tittes et al., 2021). It transforms from motile rod-shaped cells in the early exponential phase to non-motile disc shaped cells in stationary phase. Due to its aerobic growth at moderate temperatures (∼40 °C), *H. gibbonsii* LR 2-5 is further well suited for light microscopy (Tittes et al., 2021). Recently, we developed a genetic system based on deletion of the *pyrE* gene, which leads to uracil auxotrophy and allows for counter selection with 5-FOA (Tittes et al., 2024). This system enables the generation of genomic deletion mutants and the expression of genes under the tryptophan inducible promoter. In addition, the salt-stable version of GFP was shown to be functional in *H. gibbonsii* LR2-5 and allows for tagging of proteins to study their cellular localozation (Tittes et al., 2024). This combination of a broad viral susceptibility, genetic tractability and availability of molecular tools for its study makes *H. gibbonsii* LR2-5 an ideal model system for the study of euryarchaeal viruses.

HFTV1 belongs to the *Haloferuviridae*, which is part of the *Caudoviricetes*. It has a long non-contractile tail, a linear dsDNA genome, which was previously annotated so far with 68 open reading frames (Mizuno et al., 2019). Previously, we demonstrated that HFTV1 binds to the cell surface of *H. gibbonsii* within several minutes (Schwarzer et al., 2023). It uses one of the two S-layer glycoproteins, the main cell wall components, as a receptor. Binding can take place with both the capsid head and tail (Schwarzer et al., 2023). Cryo-electron microscopy has shown that the capsid head of HFTV1 is decorated with turrets, which likely bind the S-layer (Zhang et al., 2024). We hypothesized that binding with the capsid head is reversible and is followed by irreversible binding via the tail and genome delivery (Schwarzer et al., 2023; Zhang et al., 2024). HFTV1 is a lytic virus that leads to cell lysis 6 hours post-infection (p.i.). Further details of the progression of the infection cycle between the initial binding of

HFTV1 to the cell surface and cell lysis 6 hours later, are not known. We aimed to gain insight into the regulation of the genetic program behind intracellular progression of the HFTV1 infection and thus to shed light on the viral life cycle of euryarchaeal viruses. We explored the transcriptome of virus and host during infection with RNA sequencing (RNAseq). Therefore, this study provides valuable insights into the temporal regulation of the HFTV1 infection cycle and its impact on the viral host, which will pave the way for further studies on the intricacies of the HFTV1 life cycle.

## RESULT & DISCUSSION

In order to follow the progression of the HFTV1 infection, *H. gibbonsii* LR2-5 was infected and samples were taken at relevant intervals after infection (Figure 1A). To monitor the dynamics of the lytic virus life cycle, we performed virus-targeted fluorescence *in situ* hybridization (virus-targeted direct-geneFISH) using Alexa594-labelled probes specific to the HFTV1 genome (Supplemental Table S1). A progressive increase in viral signal was observed from 5 min p.i. to 300 min p.i. (Figure 1B). At early time points (5 min p.i.), single dot signals corresponding to individual viral genomes were detected, representing viral particles immediately after adsorption and injection. At 60-120 minutes after infection, an increase in fluorescence intensity was observed, indicating viral activity within the cells. The appearance of distinct fluorescent regions showed that the infection had reached a more advanced stage with active viral replication and maturation. Evidence of potential cell lysis events was observed at 5 h p.i., marked by a decline in cell population and an increase in cell debris/loss of intact cells.

**Figure 1:**
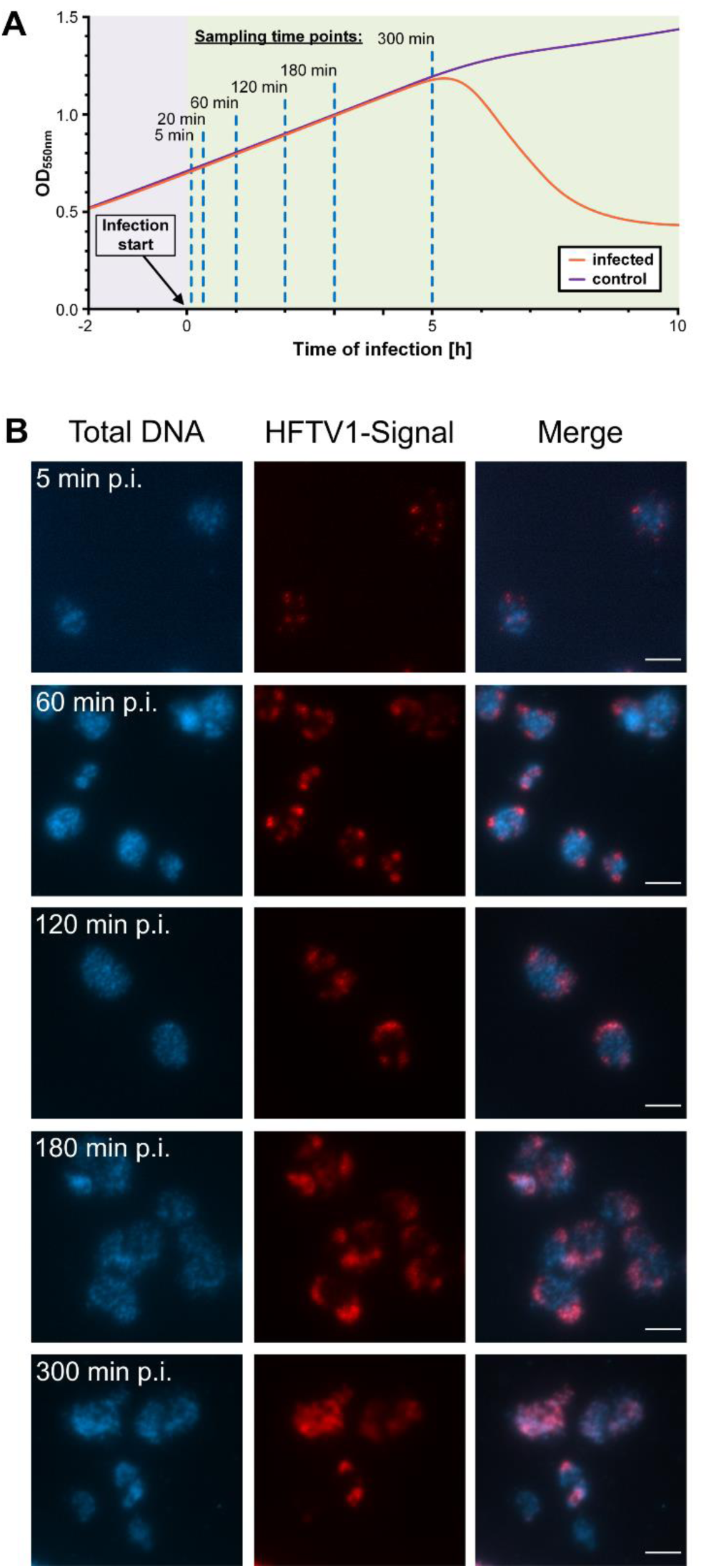
The infection cycle of HFTV1. A) Example of growth dynamics during the course of HFTV1 infection over time. Infected *H. gibbonsii* LR2-5 host cultures increase in cell abundance until lysis occurs after 5 to 6 hours, and cell densities decrease. Samples for virus-targeted direct-geneFISH and RNA isolation were collected from infected and virus-free cultures at the indicated time points. B) Fluorescence in situ hybridization of nucleic acids (virus targeted direct-geneFISH). Visualization of the infection of *H. gibbonsii* LR2-5 by HFTV1 using direct-gene FISH. First column shows the total DNA content of cells visualized by DAPI staining. The second column shows attached and intracellular virus visualized by HFTV1 targeting probes labelled with Alexa594. Third column shows the merge of DAPI and virus signal. The scale bar indicates 5 µm.

### Improved annotation of the HFTV1 genome reveals new ORFs and putative gene functions

In order to understand how the HFTV1 virus is affecting the *H. gibbonsii* LR2-5 gene expression levels and how the viral gene expression is regulated, RNA was isolated after 5, 20, 60, 120, 180, and 300 minutes p.i. cDNA libraries were prepared, subjected to Illumina sequencing and reads were respectively mapped to the viral and host genomes.

Since the original publication of the HFTV1 genome (Mizuno et al., 2019), new annotation tools and experimental data have become available (Zhang et al., 2024). Therefore, in order to improve our ORF designations in the RNAseq read mapping, we updated the annotation of the HFTV1 genome and employed Pharokka (Bouras et al., 2022) in conjunction with Phold (https://github.com/gbouras13/phold) to identify further ORFs and find predicted functions for genes of still unknown purpose.

In this process, 12 further ORFs (80 total, formerly 68) and one putative tRNA were found. In addition, some conflicts between the original and the predicted new ORF assignment were identified. In order to resolve those, we used the RNAseq data (as described below). For instance we found ORFs in the old annotation, which had start codons upstream of transcriptional start sites (TSSs), indicative that the actual start codon must be further downstream inside the transcribed region. Additionally, long untranslated regions (UTRs) pointed us to potentially overlooked ORFs. Similarly, due to the fact that unidirectional genes within the same transcript generally rely on translational coupling via termination-reinitiation (Huber et al., 2019; Huber et al., 2023; Kramer et al., 2014), we were confident that newly annotated genes within intergenic regions of a continuous transcript, are likely to be correct. The resulting curated annotation is shown in Figure 2 and was used for the alignment of the following HFTV1 RNAseq analysis.

**Figure 2.**
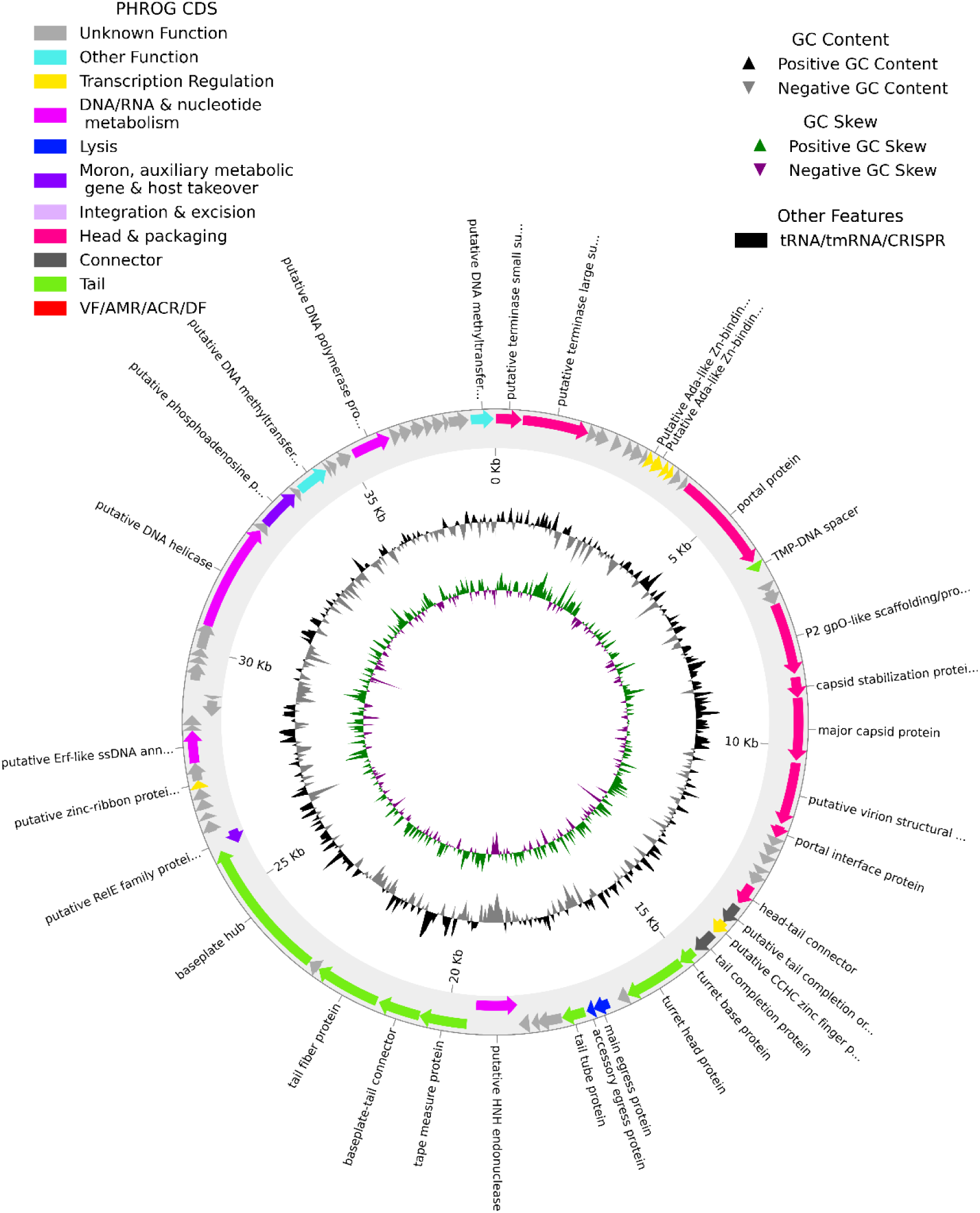
Updated annotation of the HFTV1 genome. The reference genome (NC_062739.1) was updated with pharokka and phold (see M&M) as well as with RNAseq. Different colours indicate genes of different predicted functional groups, based on their respective PHROG (Prokaryotic Virus Remote Homologous Group) assignment.

### Expression of viral genes

To gain insight into the differential expression of HFTV1 throughout the infection, the HFTV1 reads were normalised, for the total amount of reads (HFTV1 and *H. gibbonsii* LR2-5 combined) and gene length using SeqMonk V1.48.1. Due to the short incubation time at timepoints 5 and 20 minutes p.i., only a small fraction of all reads align to the HFTV1 viral genome. However, this number of viral reads rapidly increases over time. Based on the temporal trend of reads mapping to annotated genes (and one tRNA) we divided the HFTV1 genes in ‘early’ (coloured red), ‘middle’ (coloured green), ‘middle-down’ (coloured black), and ‘late’ (coloured blue) expression groups, (Figure 3A). For this classification we assessed the expression per gene per timepoint by normalizing the read coverage of genes by their length and total reads, followed by calculating a percentage coverage relative to the respective maximum coverage for each gene (over all five timepoints). As shown in Figure 3B, the grouping was then assigned whether genes reached their half-maximum expression (dotted line) by 20 minutes (“early”; 6/81; 7% total), 60 minutes (“middle” 20/81; 25% total) or after 60 minutes (“late” 44/81; 54% total). Further, genes that reached their half-maximum expression by 60 minutes, but later again fell below this half-maximum threshold are referred to “middle-down” genes (11/81; 14% total). The relative percentage coverage was also plotted over time for each gene of the respective temporal categories showing the clustering into expression trends (Figure 3 C-F)

**Figure 3.**
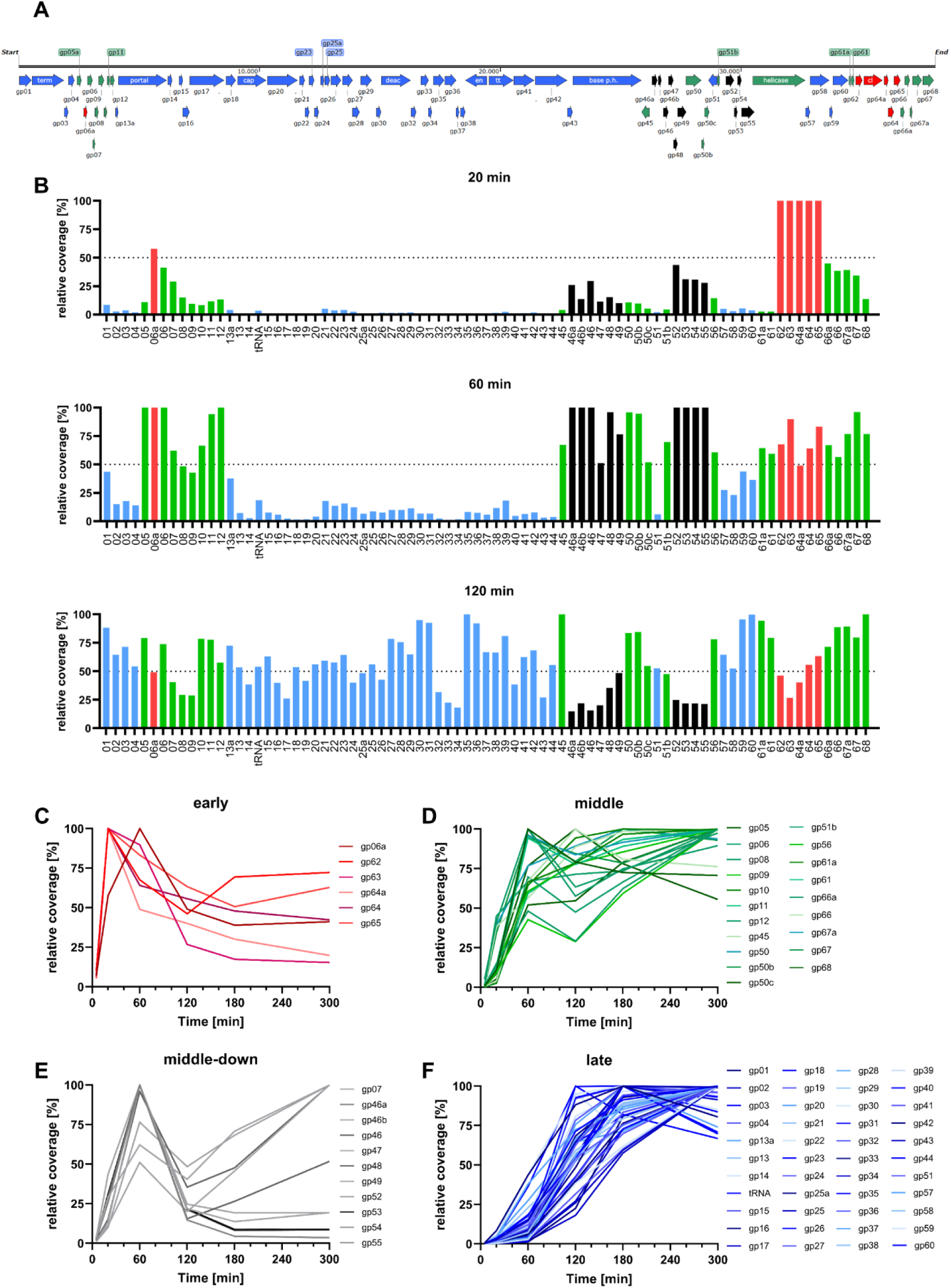
Expression of HFTV1 genes during viral infection of *H. gibbonsii* LR2-5. (A) Schematic representation of the newly annotated HFTV1 genome. (B) Gene length normalised gene expression profiles of HFTV1 at 20, 60, and 120 minutes p.i. The y-axis represents the percentage of the total gene expression of the individual genes. The early, middle-down, middle and late expressed gene groups are represented in red, black, green and blue, respectively. The total percentage coverages of (length normalized) viral ORFs are shown over time, for genes clustering into expression groups introduced in panel (B), namely: (C) early, (D) middle-down, (E) middle, and (F) late. The results shown in panel B-F reflect the averages of three biological replicates. *Gp62*-*gp65* are all expressed at an ‘early’ onset with *gp63* (PCNA, DNA polymerase sliding clamp) being the only annotated gene. This is most likely required for viral DNA amplification, known as one of the first steps in the viral infection cycle (Mizuno et al., 2019). Additionally, many structural proteins, or proteins required for the assembly of the virus particle, are expressed in the late phase of the viral infection cycle including: *gp13* (portal protein), *gp17* (P2 gpO-like scaffolding/protease protein), *gp18* (capsid stabilization protein), *gp19* (HK97 gp5-like major capsid protein), *gp29* (gp17-like tail completion protein), *gp35* (tail tube protein) and *gp40* (phage tail tape measure protein).

A noteworthy feature of the HFTV1 expression is that some gene clusters that appear like *bona fide* operons, are highly differentially expressed. For example, gene cluster *gp52 - gp60* appears like a *bona fide* operon, based on its genomic lay out (Supplemental Figure 1A). G*p52 - gp60* seem to be located after a common promoter and the genes have overlapping stop and start codons without intergenic sequences. However, the trend in their expression pattern over time is significantly different for the various genes located in the same cluster. In the gene cluster *gp52 – gp60, gp52 – gp55* are ‘middle-down’ genes, indicating that their onset of expression is in the middle of the infection cycle and subsequently is decreased severely (Figure 3D), and *gp56-60* are “middle” to “late” genes, which are not downregulated in late stages of infection (Supplemental Figure 1B). This pattern is indicative of intergenic promoters, which in *H. volcanii* have been shown to account for ca. 6% of all identified transcripts (Laass et al., 2019). Therefore, similarly we manually annotated all transcriptional start sites (TSSs) in the RNAseq mapping, based on the typical sudden spike in coverage at exactly the same position in the genome. Indeed, during this process we discovered an intergenic promoter within gp 56 (TSS at 31.839 nt), which explains why the genes of the supposed gp52-60 operon appear to follow two distinct temporal expression trends. Interestingly, based on coverage distribution 56 appears to be a middle gene. However, since the intergenic promoter is located within the ORF, the RNA coverage stemming from this internal TSS does not contribute to translatable gp56 mRNA. This finding effectively places gp56 in the middle-down (almost early) gp52 to 55 group, which due to its hypothetical function in genome replication also fits conceptionally. This suggests that the two promoters of the gp52-60 region are differently regulated, which may be evidence of the high level of optimization in genome organization that has taken place in this virus to allow for maximum coding capacity while limiting genome size expansion.

### Organisation of the viral genome reveals clear antisense regulation of functional groups

As the transcriptional organization of the HFTV1 genome appears to be fairly complex and intergenic promoters may lead to incorrect assignment of the temporal expression group of genes (i.e. gp56), we proceeded to organize the HFTV1 ORFs into operons. We assigned genes into the same operon based on the following three criteria: (***i***) neighboring start-stop codons, (***ii***) clustering into the same temporal expression profiles (cfr. Figure 3, with the exception of intergenic promoters) and (***iii***) a continuous coverage throughout the proposed transcript (Figure 4A).

**Figure 4:**
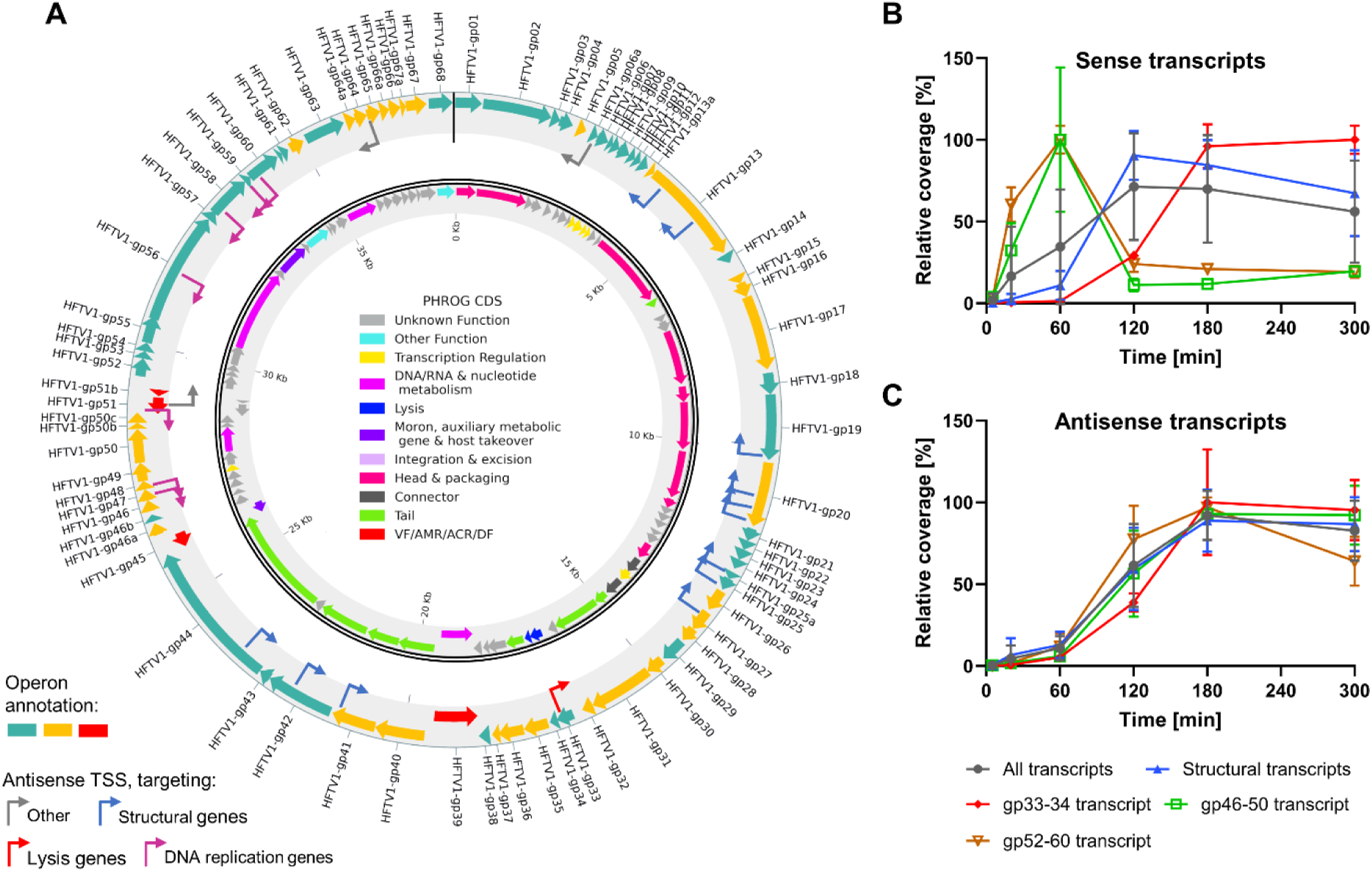
Transcriptional organization of the HFTV1 genome. A) Outer ring: Assignment of the HFTV1 ORFs into operon structures, indicated with alternating coloration of ochre and teal for forward operons and red for reverse operons. The bent arrows indicate the identified transcriptional start sites (TSS) and are colored according to the types of genes which are in the transcripts that the corresponding antisense RNA would hybridize with. Inner ring: Functional grouping of the HFTV1 genome (cfr. Figure 2). B) Relative percentage coverage over time shown of selected viral sense transcripts associated with antisense RNA interactions during the HFTV1 infective life cycle. C) Relative percentage coverage over time shown of viral antisense transcripts during the HFTV1 infective life cycle. For B and C error bars represent the averages and standard deviation of the coverages of multiple TSS, or, in case data of a single TSS is shown, averages and standard deviations among the tree replicates for the same data point.

As a result, we were able to cluster the 80 thus far identified ORFs into 28 distinctly regulated transcriptional units. Of these, 12 are monocistronic (42.8%), while the overall HFTV1 genome has an average of about three genes per operon. This is in stark contrast to phages, which generally have few but highly polycistronic operons (Jack et al., 2019; Kornienko et al., 2020; Oppenheim et al., 2005; Putzeys et al., 2023). Therefore, the transcriptional organisation of HFTV1 more closely mirrors that of its host. In *Haloferax volcanii* about 2/3 of the transcripts are monocistronic and thus each operon on average has thus a relatively small number of genes (1.7) (Laass et al., 2019). As such, the HFTV1 operon structure is intermediate between that of canonical phages and eukaryotic viruses, which are one of few instances where polycistronic operons can be found in this transcriptionally fundamentally different domain (Blumenthal, 2004; Shine et al., 2024).

Curiously, during this process of assignment of HFTV1 reads to transcriptional units, we observed for certain operons that many reads also mapped to the reverse strand. Therefore, we also identified the TSSs of these putative antisense RNA (asRNA) transcripts, which have been shown to play important regulatory roles in *Haloferax* species (Babski et al., 2016; Gelsinger & DiRuggiero, 2018; Laass et al., 2019).

As shown in Figure 4A, more than half (52.25%, 12/23) of these potential antisense transcripts (indicated as bend arrows) are located in or at the end of operons associated with structural genes for the formation and eventual packaging of new capsids. A second apparent hotspot for asRNA (30.48%, 7/23) are two operons containing genes associated with DNA replication. In order to get a better understanding of potential mechanisms behind these observations we quantified the transcription profiles of all identified promoters in the HFTV1 genome based on the relative coverage in their respective TSS over time, and compared sense transcripts with their antisense counterparts (Figure 4BC, Supplemental Table S2). The average of all sense transcripts (Figure 4A grey line) does not show any temporal trend, which is indicative of genes with different (i.e. early or late) temporal expression profiles averaging out the coverage change over time. In contrast, we found that the average of all antisense transcripts followed a “late” expression profile, peaking at around 180 minutes.

The sense transcripts mapping to genes of different functionalities, show a very clear temporal pattern of early mid and late genes. We can identify typical early functions such as DNA metabolism (gp46-50 and gp52-60) peaking at 60 min p.i., structural genes that are highly expressed only from 120 min (mid genes) and finally the proposed (late) lysis transcript (gp33-34, Kuiper personal communication), which is only highly expressed from 180 minutes and reaching its maximum at 300 minutes, when host lysis occurs.

Therefore, the expression profiles of the identified HFTV1 transcripts appear to follow the typical viral life cycle progression patterns from genome replication ◊ to virion production ◊ to egress (Birge, 2006; Desiere et al., 2000; Lawrence et al., 2009; Liu et al., 2013; Ortmann et al., 2008; Ventura et al., 2002). Interestingly though, no alternative RNA polymerase has been identified in the HFTV1 genome and no transcript elongation at later time points is evident (suggestive of i.e. antitermination) (Figure 3B). Therefore, it appears that the regulation in the HFTV1 life cycle is accomplished by alternative mechanisms to regulate time-resolved transcription initiation regulation in conjunction with asRNA expression.

Occasionally asRNA can also be found in phages, where it effectively silences the translation of the sense transcript (i.e pOOP or paQ asRNA) (Oppenheim et al., 2005; Owen et al., 2020). Therefore, the late expression of nearly all asRNAs (such as those targeting the operons gp46-gp50 and gp52-gp60), suggests that they play an important role in down regulating the transcripts of genes involved in early infection functions such as DNA metabolism. This is likely to shift all resources to particle maturation and egress.

Generally, it seems that asRNA based regulation is much more involved in the life cycle regulation of HFTV1 than previously anticipated. However, this might also be based on methodical biases, as many studies on phages pre-date the availability of RNAseq and thus may have overlooked asRNAs. For instance, for the Salmonella phage BTP1 recently new asRNAs have been identified via RNAseq (Owen et al., 2020). Interestingly, one of these asRNAs (STnc6030), which is expressed during lysogeny appears to mediate superinfection immunity by silencing a structural transcript, while still permitting prophage induction (Owen et al., 2020).

### Impact of viral infection on host gene expression

Next, we analyzed how viral infection affects host gene expression. An intensity difference statistical analysis of *H. gibbonsii* LR2-5 infected with HFTV1 showed 0, 7, 135, 247, 434 and 374 differentially expressed host genes after 5, 20, 60, 120, 180 and 300 minutes p.i. compared with the uninfected control culture, respectively. At the earlier timepoints (20 and 60 minutes) the majority (100 and 88%, respectively) of the differentially expressed genes are up regulated whereas later in the infection cycle a more equal distribution between up and down regulated genes is observed. Figure S2 illustrates the overlap and specificity of upregulated (Figure S2A) and downregulated (Figure S2B) DEGs at 5, 20, 60, 120 and 300 minutes p.i.. While the majority of DEGs were up- or downregulated specific to individual time points, a small core set of upregulated genes (e.g. 33 genes; Figure S2A) was shared across all time points starting from 60 minutes p.i.. Similarly, a subset of 39 genes was consistently downregulated from 120 minutes onwards. These patterns suggest a (possibly also virus mediated) host response to the viral infection by differentially regulating genes potentially involved in antiviral responses and, especially for mid-to-late infection stages, an apparent suppression of specific pathways in a coordinated manner.

Subsequently, we aimed to identify groups of functionally related genes that are significantly overrepresented among up or down regulated genes. EggNOGv5.0 was used to classify all differentially expressed *H. gibbonsii* LR2-5 genes in arCOGs. Using the genome of *H. gibbonsii* LR2-5 as template, 2321 genes were successfully placed in a functional category whereas of the remaining 1731 genes (42.7 %), the function was listed as unknown (arCOG category S). All differentially expressed genes, divided into up- and down-regulated genes, were linked to their specific arCOG category. The arCOG analysis for the 5, 20 and 60 minutes p.i. time points was omitted due to the too low number of differentially expressed genes, which distorts the analysis (Figure 5).

**Figure 5.**
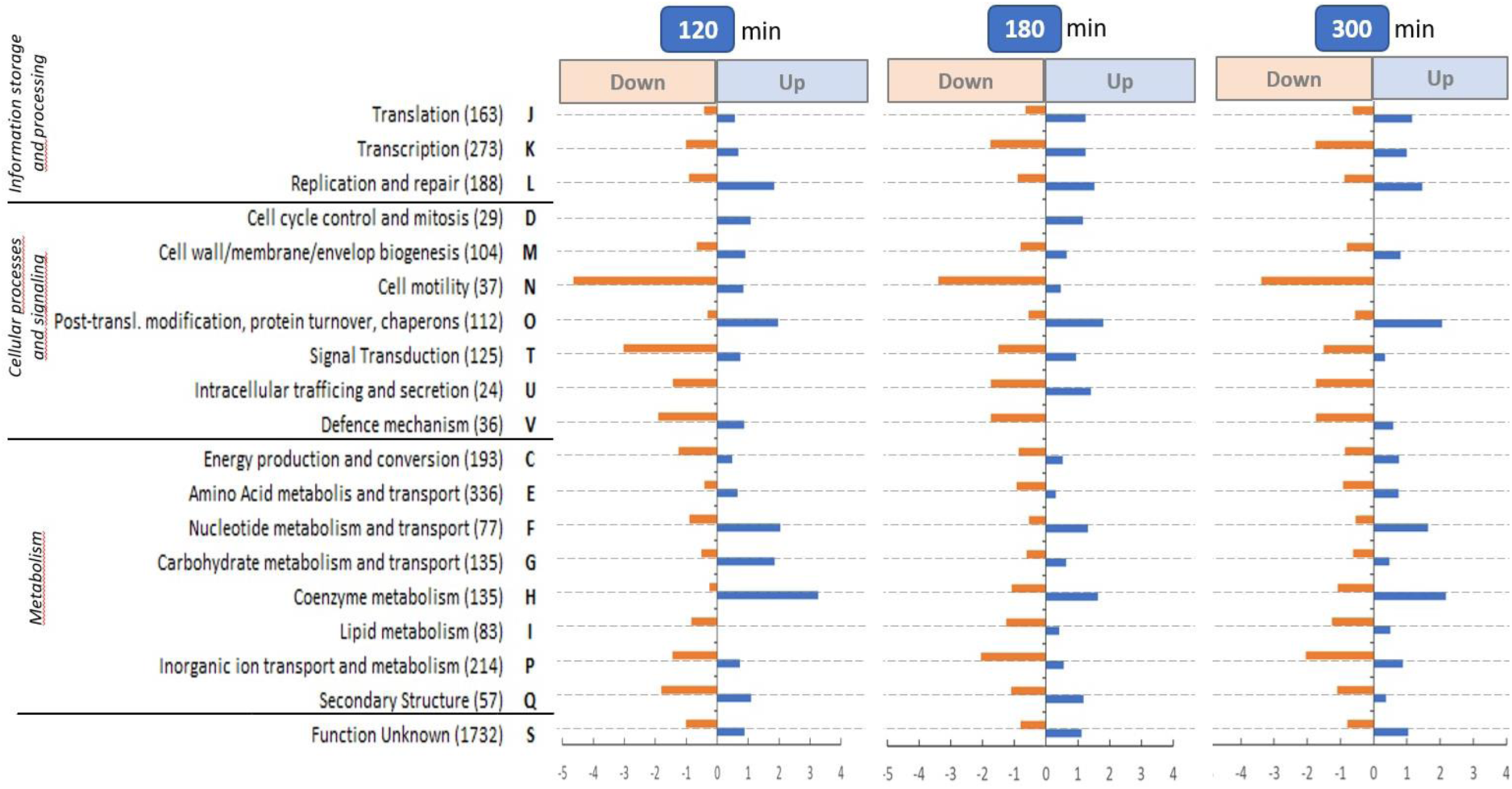
arCOG gene set overrepresentation analysis of differentially expressed down-regulated (in orange) and up-regulated (in blue) genes of *H. gibbonsii* LR2-5 upon infection of HFTV1 after 120, 180 and 300 minutes p.i., All genes were categorized in arCOGs and clustered in 4 groups: information storage and processing, cellular processed and signalling, metabolism and unknown function. The numbers in parentheses indicate the total number of genes in the arCOGs. The over (or under) representation was calculated based on a random distribution of all differentially expressed genes over all arCOGs. *E.g.* in the cell motility arCOG (N) there are 37 genes. The *H. gibbonsii* LR2-5 genome contains 4053 genes therefore 0.91% of all genes belong to arCOG N. At T120 118 genes are downregulated of which 5 belong to arCOG N, which is 4.24%; whereas 0.91% is expected. Therefore arCOG N is 4.64 times over-represented of the down-regulated genes at T120 (See x-axis left panel; Cell motility (37) N). A value of 1 is expected and > 1 means an over-representation and < −1 and under-representation of the number of differently expressed genes in that arCOG.

### Increased expression of genes involved in nucleotide metabolism

Several genes belonging to ArCOG F, ‘Nucleotide metabolism and transport’, were strongly upregulated in infected cells, after 120, 180 and 300 minutes p.i. compared to uninfected controls (Figure 5). This likely reflects the increased DNA replication and transcription activities in order to replicate the HFTV1 genome. Several of the genes belonging to this group are amongst the highest upregulated genes 120, 180 and 300 min p.i.. For example, several genes involved in pyrimidine synthesis such as *dcd* (a dCTP deaminase, (Johansson et al., 2003)), *pyrG* (which catalyzes the ATP-dependent amination of UTP to CTP), *guaAb* (which catalyzes the synthesis of GMP from XMP) (Jewett et al., 2009), and *uraA* (a xanthine/uracil permease) are all ∼5 × upregulated at 120 and 180 min p.i.

Dcd in archaea is responsible for the conversion of dCTP to dUTP and thus can help to adjust levels of dCTP in relation to dTTP in the nucleotide pool (Huffman et al., 2003). In a next step, the dUTP can be acted upon by dUTPase to form dUMP, in order to prevent incorporation of dUMP in the DNA (Huffman et al., 2003). In bacteria, PyrG is a CTP synthetase, which catalyses the glutamine- and ATP-dependent amination of UTP to form CTP and its expression is regulated independently of the rest of the *pyr* operon containing pyrimidine biosynthetic genes (Jewett et al., 2009). The role of this gene has not been studied in archaea. The UraA protein involved in uracil/xanthine transport has been described to be highly transcribed in *Ca*. *Thalassarchaeaceae,* but further characterization of this protein in archaea is lacking (Damashek et al., 2021). NrdJ, the adenosylcobalamin (Vitamine B12)-dependent ribonucleoside-diphosphate reductase, catalyzes the conversion of ribonucleotides to deoxyribonucleotides which is essential for de novo DNA synthesis (Borovok et al., 2006; Eklund et al., 2001; Rodionov et al., 2003) and this gene is also one of the members of ArCOG F; ‘Nucleotide metabolism and transport’. NrdJ is, in the uninfected control condition, one of the highest expressed genes in *H. gibbonsii* LR2-5 and is, after HFTV1 infection, upregulated 2.0, 2.1 and 2.0 times after 120, 180 and 300 minutes p.i., respectively. Like many resources required for the amplification of viral particles, NrdJ most likely is upregulated due to the high demand for nucleotides in order to replicate the viral genome.

### Upregulation of post-translational modification and chaperonins

The arCOG O, “Post-translational modification, protein turnover, chaperone functions”, was also amongst the enriched arCOGs at 120, 180 and 300 minutes p.i.. Upregulated genes belonging to this class, are encoding several different chaperonins (such as the TCP1 and DNA-J class). Also, a membrane bound mannosyl transferase, encoded by HfgLR_07500, is significantly upregulated in infected cells. This protein is likely involved in addition of the N-linked glycan biosynthesis, required for N-glycosylation of archaeal surface proteins (Gandini et al., 2020). Upregulation of this class of proteins (arCOG O) could be explained by the high demand of protein expression, folding and post-translational modification machinery due to the massive expression of HFTV1 proteins in the hours after infection leading up to production of mature viral particles.

### Upregulation of the coenzyme metabolism by precorrin pathway

At 120, 180 and 300 minutes p.i., amongst all upregulated genes, the “coenzyme metabolism” arCOG (H) was 2.4, 3.3 and 1.6 times over-represented, respectively, (Figure 6). ArCOG H contains (in total) 135 genes of which, at *e.g.* 300 minutes p.i., 13 are significantly upregulated (Figure 6). Ten out of the 13 upregulated arCOG H genes (annotated as cbi. And cobN) belong to the precorrin pathway which produces precorrin-2, the precursor of cobalamin (or vitamin B12).

**Figure 6.**
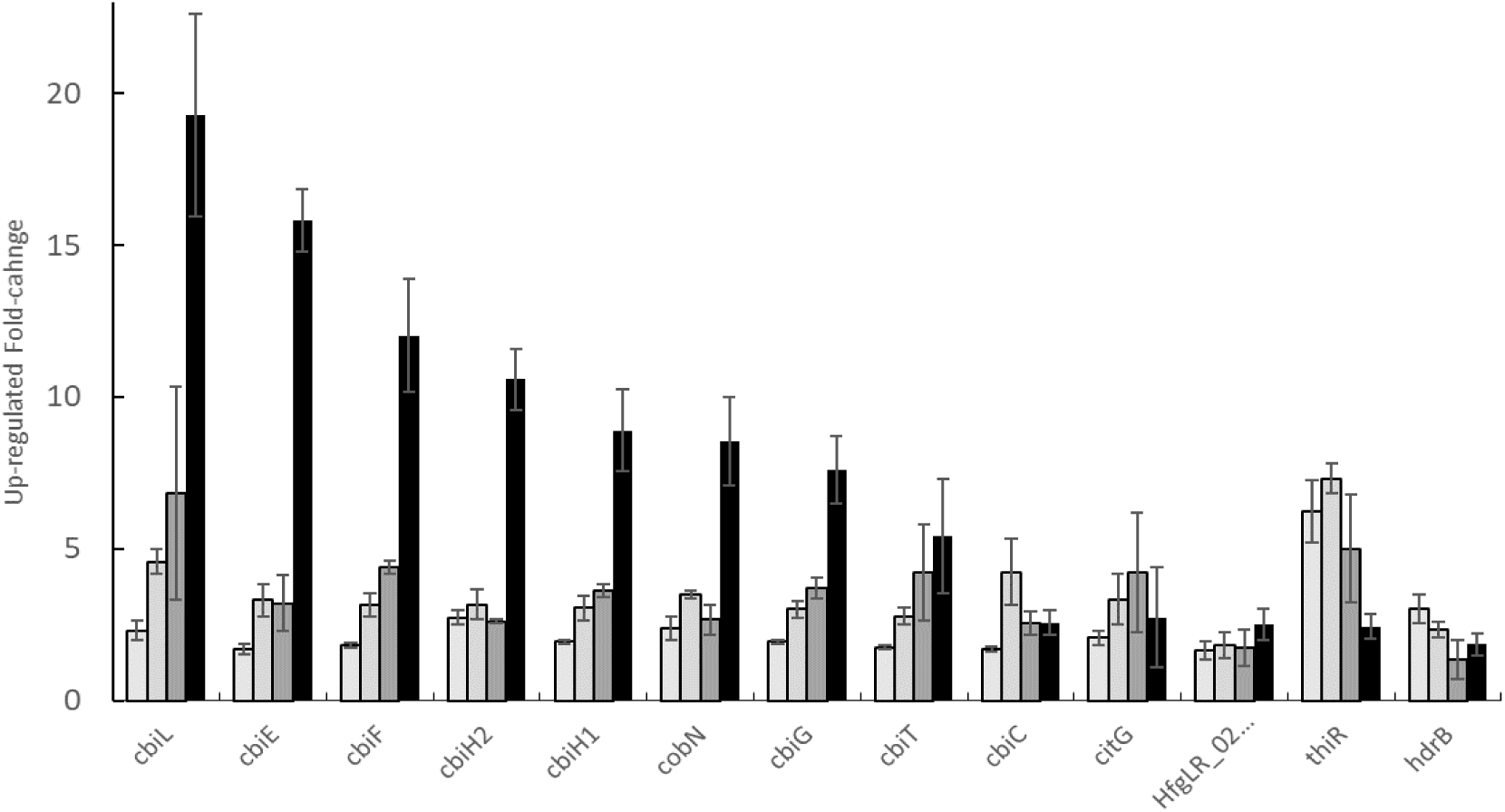
Fold change of the major up-regulated genes of the “Coenzyme metabolism” arCOG H at 60 (¢), 120 (¢), 180 (¢), and 300 (′), minutes p.i.. Error bars indicate the standard deviation of three biological duplicates.

In bacteria and archaea vitamin B12 is required in various enzymes which include methionine synthase (Banerjee & Matthews, 1990), ribonucleotide reductase (Borovok et al., 2006; Eklund et al., 2001; Rodionov et al., 2003), glutamate and methylmalonyl-CoA mutases (Takahashi-Iñiguez et al., 2012), ethanolamine ammonia-lyase (Wetmore et al., 2002), and diol dehydratase (Toraya & Fukui, 1977). We hypothesize that the upregulation of the precorrin pathway is a demand driven need for vitamin B12 which is required for *e.g.* NrdJ, the Vitamine B12-dependent ribonucleoside-diphosphate reductase, see above.

### Cell motility genes are downregulated

Amongst the down-regulated genes, the arCOG category N, “cell motility”, was significantly overrepresented with 4.6, 3.4 and 3.4 times more representatives than expected at 120, 180 and 300 minutes p.i, respectively (Figure 5). Belonging to this arCOG “cell motility” are genes encoding adhesive pilins. Specifically, *pilB4* and *pilC4* showed significantly reduced expression in infected cells. PilB and PilC are an ATPase and membrane anchor, respectively, that make up the motor of adhesive type IV pili in archaea (Esquivel et al., 2013). The pilins that make up the filament are encoded by *pilA* and loaded into the filament by the PilB/C complex. These pili are homologous to bacterial type IV pili and are involved in adhesion and biofilm formation (Esquivel et al., 2013). Haloarchaeal genomes usually contain multiple different copies of *pilA*, *pilB* and *pilC,* for instance the *H. volcanii* genome encodes for five PilB/PilC pairs.

*H. gibbonsii* LR2-5 encodes 4 PilB-PilC pairs and 6 different PilAs. All of these remain stable throughout the infection, with the exception of PilB4-PilC4. PilB4-PilC4 is homologous to the same pair from *H. gibbonsii*. The expression of the PilB4-PilC4 pair is under normal uninfected conditions higher than those of PilB3-PilC3. Downregulation of PilB4-PilBC4 leads to levels that are only lower than that of PilB3-PilC3. Thus, downregulation of PilB4-PilC4 could lead to less piliated cells or to cells with pili that have a different PilA composition. As the pili from *H. gibbonsii* were not yet analyzed in detail, the impact of PilB4-PilC4 remains to be established.

Five significantly down-regulated genes that are part of the ”cell motility” category are *arlA1, arlA2, pilB4, pilC4* and *cheW1* which show a fold change of 0.44, 0.29, 0.38, 0.44 and 0.43 at 120 minutes p.i., respectively. ArlA1 and A2 are the main archaellum components of *H. gibbonsii* that form the filament of this rotating motility structure (Tittes et al., 2021). Besides downregulation of the archaellins *arlA1* and *arlA2*, almost all archaellum genes in this operon were downregulated starting at 120 minutes p.i. (Figure 7). The other *arl* genes, *arlG, arlJ, arlF* and *arlCE*, in the operon, encoding different components of the archaellum motor, show a fold change, a of 0.43, 0.52, 0.76 and 0.83, respectively. Thus, HFTV1 infection leads to clear and strong downregulation of the archaellum machinery.

**Figure 7.**
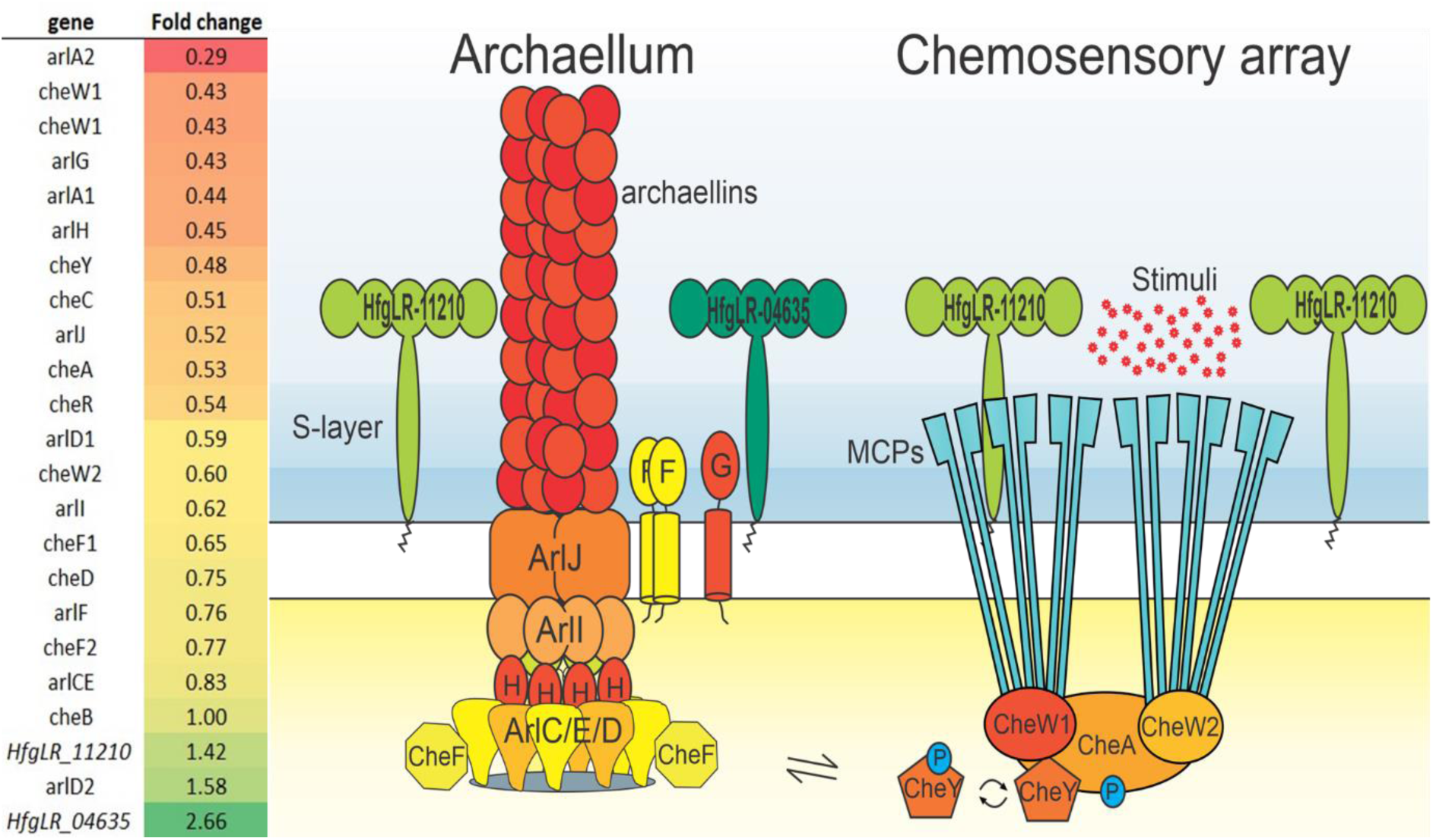
Fold change values of a subset of genes of the archaellum and chemosensory array of the “cell motility” arCOG N at 120 minutes p.i. in which the colours used in the image match the colour scheme of the fold change values indicated on the left. HfgLR-11210 and HfgLR-04635 are the S-layer encoding genes.

CheW1 is a component of the chemosensory arrays, together with CheA, the response regulator CheY and the MCPs (methyl-accepting chemotaxis proteins). These latter detect extracellular signals and transfer them via the chemosensory arrays to the archaellum, in order to respond adequately to environmental ques. Besides *cheW1*, also *cheC, cheR, cheW2* and *cheD*, all encoding components of the chemotaxis system, were 1.9, 1.8, 1.7 and 1.3 times downregulated, respectively, at 120 minutes p.i.. Their expression decreased even further at 180 and 300 minutes p.i.. (Figure 7). Another gene in the top 10 of downregulated genes at 120 min p.i, is *hfgLR_13260*, which is annotated as a PAS domain containing histidine kinase. PAS domain containing proteins are known to sense environmental factors, such as light, nitrogen, oxygen or redox potential. These sensing proteins are usually signaling downstream systems, such as the motility machinery. For example, the methyl-accepting chemotaxis proteins that are the sensory receptors part of the chemotaxis system (see below) can contain such PAS domains. It remains to be established if *hfgLR_13260* is transferring signals to the motility machinery.

Together, these results suggest that the expression of the large part of the chemotaxis machinery is downregulated after HFTV1 infection. Reduced expression of both the archaellum and the chemotaxis system, suggest that HFTV1 infected cells become less motile after infection.

### Cells show reduced swimming behavior after infection

Time lapse microscopy was applied to analyze if the downregulation of the archaellum and chemotaxis operons would lead to obvious reduction of swimming behavior of infected cells. Cells were grown to mid-exponential phase, infected with an M.O.I. of 10 and compared with an uninfected control strain grown under similar conditions at different time points p.i.. Cells were visualized by time lapse microscopy under native growth temperature. Time lapse movies were made and cells were tracked (Supplemental Video 1 and Supplemental Video 2). The average velocity of cells was measured in each field of view (Figure 8). Indicating that HFTV1 infected cells had a greatly reduced tracking velocity compared to healthy control cells. Whereas control cells swim with an average velocity of 5.4 µm/s, 1 hour after HFTV1 infection, the velocity was already reduced to 2.2 µm/s and further reduced to 0.4 µm/s at 5 hours post infection.

**Figure 8:**
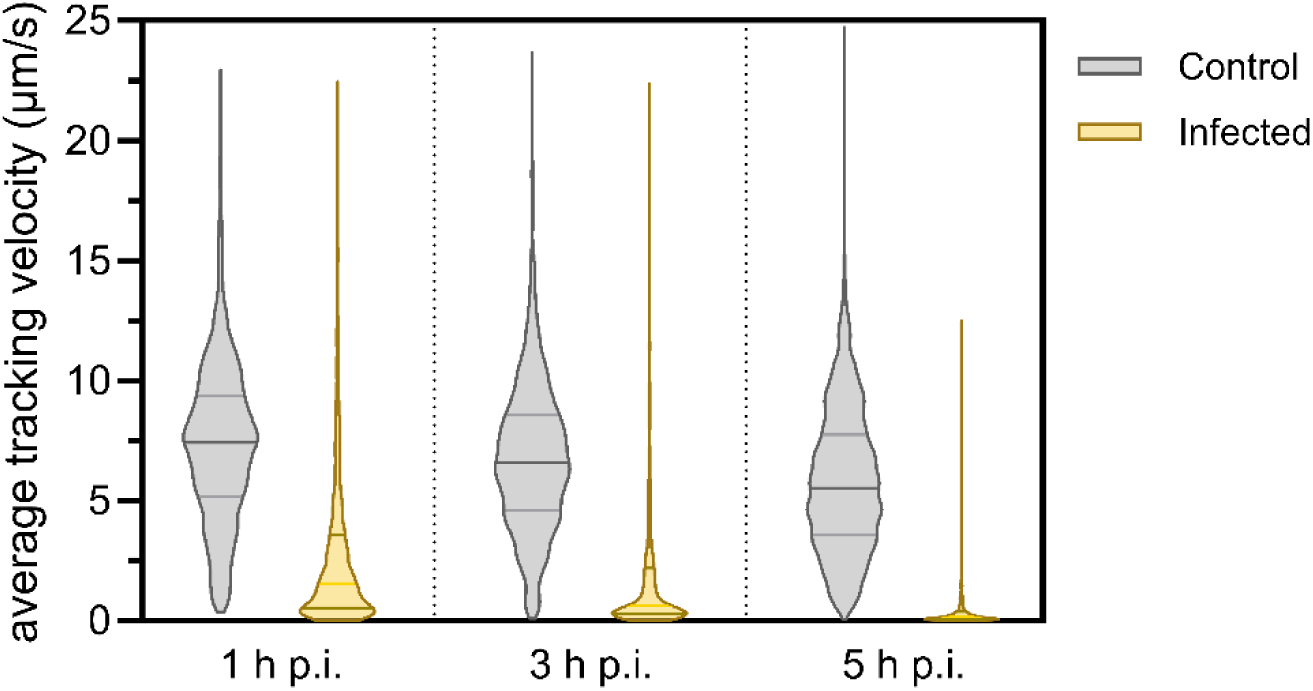
Infection of HFTV1 inhibits swimming motility of *H. gibbonsii*. Violin plots depict the distribution of mean tracking velocities (µm/s) for cells in control (grey) and HFTV1-infected (yellow) cultures at 1, 3, and 5 h p.i. Tracking and quantification were performed using TrackMate. Average tracking velocity was measured over 15 sec for each indicated condition and time point. HFTV1-infected cells exhibited a reduced mean tracking velocity compared to uninfected controls, throughout the infection cycle, indicating virus-induced impairment of cellular motility. Sample sizes were as follows: 1 h p.i., n = 1296; 3 h p.i., n = 973; 5 h p.i., n = 1705. Statistical analysis was conducted using unpaired Student’s t-tests comparing infected and control groups at each time point. All comparisons yielded highly significant differences (p < 0.0001).

Thus, HFTV1 infected *H. gibbonsii* LR2-5 cells are significantly less motile compared to non-infected controls. The reduced motility is likely due to the specific downregulation of the archaellum operon. This could be a consequence of redistribution of energy for the replication and production of HFTV1 particles. Alternatively, rendering their hosts non-motile, might somehow provide HFTV1 with an advantage, for example by promoting a sessile life style in biofilm, where the availability of potential new hosts will be higher.

Inhibition of expression of motility genes has been reported to occur after infection by several bacteriophages. For example, motility genes are repressed early in the infection cycle of bacteriophage PaP3, infecting *Pseudomonas aeruginosa*, the filamentous SW1 phage infecting the bathypelagic bacterium *Shewanella piezotolerans* and the filamentous ϕAFP1 phage, infecting the gram-negative marine bacterium *Alteromonas abrolhosensis* (Jia et al., 2022; Jian et al., 2013; Jian et al., 2016; Zhao et al., 2016). It is generally assumed that these viruses inhibit expression of motility genes in order to balance the high energy demand associated with viral replication and particle production. And this is likely also occurring after HFTV1 infection together with the above mentioned manipulation in host metabolism and chaperonis expression. Virus-beneficial changes in the transcriptional landscape of host during infection have also recently been observed for halovirus His2 infecting *Haloarcula hispanica* (Lee et al., 2020), as well as in some, but not all viruses infecting hyperthermophilic archaea, and as such seems to be an important aspect in the reproductive strategy of archaeal viruses (Fröls et al., 2007; Okutan et al., 2013; Ortmann et al., 2008; Ren et al., 2013).

## CONCLUSION

In conclusion, the here presented data provides a detailed picture of transcriptional regulation during the infective life cycle of the model archaeal virus HFTV1. We show that HFTV1 infection significantly impacts host gene expression. The observed increased expression of genes involved in nucleotide metabolism and chaperonis, is likely orchestrated to support viral replication and assembly of viral particles. We also observed a specific high upregulation of the precorrin pathway responsible for vitamin B12 production, needed for B12 dependent enzymes. Lastly, upon infection, motility and chemotaxis genes specifically are quickly downregulated, which is in correspondence with reduced swimming speeds of infected cells, as observed by time lapse microscopy. Cellular motility could be actively repressed by the virus to redirect resources from energy-costly motility towards other process, or it might be linked with specific benefits for the viruses in sessile cells, such as a potentially reduced likelihood for subsequent encounters with viruses. Our analysis of viral transcripts on the other hand shows that HFVT1 expression is tightly regulated throughout the infective life cycle, similarly to what is typically observed for bacterial viruses. Interestingly though, we further observed fine-tuned temporal transcriptional initiation from within gene clusters, which can provide HFTV1 with extra opportunities for individual gene regulation.

Additionally, the high number of transcripts vs number of predicted genes in HFTV1, is in stark contrasts with bacterial viruses, where generally many genes are arranged in a few long transcripts. Another, striking difference with bacterial viruses, is the prominent role of asRNA in the regulation of late-expressed genes of HFTV1. The observed important role of asRNA in this archaeal viral model, might be linked to the high presence of asRNA in archaea compared to bacteria (Laass et al., 2019).

Generally, in comparison to temporal transcriptomic analyses that have been performed on other archaeal viruses, the complexity of the large number of differentially regulated sense and antisense transcripts in HFTV1 appears to stand out. However, so far almost all other archaeal viruses for which time resolved transcriptomic data exists are hyperthermophiles and/or viruses with a significantly smaller genome (<20kb) (Fröls et al., 2007; Gehlert et al., 2022; Kessler et al., 2004; Lee et al., 2020; Ren et al., 2013). The only other halovirus for which time resolved microarray data has been published is the chronic virus His2, which with a genome of 16 kb encoding 35 ORFs shows clear temporal regulation, but only has three distinctly regulated regions (i.e. early, middle and late; (Lee et al., 2020)).

This work provides unprecedented insight into transcriptional regulation of haloarchaeal viruses, which will be valuable for future studies of archaeal transcription and increases our insight into the mechanisms by which archaeal viruses take control of their host cells.

## MATERIALS AND METHODS

### Growth and infection

Growth and infection procedures were carried out as previously described (Schwarzer et al., 2023)., with detailed conditions provided in the Supplementary Text.

### Probe design, synthesis and chemical labeling

Polynucleotide probe design, synthesis, labeling, and specificity testing were performed according to established protocols (Moraru, 2021), with probe sequences and labeling details listed in Supplemental Table 1 and Supplementary Text.

### Virus targeting direct-geneFISH (virusFISH)

VirusFISH was conducted following the direct-geneFISH protocol by Barrero-Canosa and Moraru (2021), with modifications described in the Supplementary Text.

### Bioinformatic genome analysis

Genome annotation and analysis were conducted using Pharokka and associated tools as described in Supplementary Text.

### Sample preparation and RNA isolation

Flash frozen cell pellets were thawed on ice and total RNA was extracted using RNeasy® Plus Mini Kit and RNeasy MinElute® Cleanup Kit (Quiagen, Hilden, Germany) according to the manufacturer’s instructions, following protocol 1 for purification of total RNA containing miRNA. RNA concentration were measured by spectrophotometer using a Nanodrop (Implen, Munich, Germany).

### Expression analysis

RNA-seq and differential expression analyses were performed by Vertis Biotechnologie AG and analyzed using standard pipelines (BowTie2, DESeq2, SeqMonk), as described in Supplementary Text.

### Swimming analysis with time-lapse microscopy

Swimming behavior was assessed using time-lapse phase-contrast microscopy at 45 °C, as detailed in the Supplementary Text.

## Data availability

FastQ files from RNA sequencing, were deposited at Zenodo and can be accessed via the following DOI: 10.5281/zenodo.15221547. Any code used for data processing and appropriate documentation can be found in Materials & Methods.

## Author contributions

Conceptualization of the work, T.E.F.Q; Writing original draft S.S., L.E.B., J.G.N. and T.E.F.Q, Writing review and editing, all authors; VirusFISH, S.S., I.H.A., and C.M.; Acquisition, analysis and interpretation of microscopy, S.S., C.M.; Viral genome annotation, L.E.B.; Investigation and validation of RNA-Seq data, S.S., L.E.B., J.G.N., T.E.F.Q, A.J., C.S.; Funding acquisition, T.E.F.Q. and L.E.B.; All authors have read and agreed to the published version of the manuscript.

## Conflict of interest

The authors declare no conflict of interest.

## Acknowledgements

This work was supported by the Deutsche Forschungsgemeinschaft (German Research Foundation) with an Emmy Nöther grant (411069969). Further financial support is acknowledged from an ERC starting grant (101039446 ARCVIR) and a Vidi grant (Vidi.223.020) from the Netherlands Research Council (NWO) to T.E.F.Q. This work was additionally supported by a Human Frontiers Science Program Young Investigator Grant (RGEC33/2023) and an interdisciplinary grant from the Hector Fellow Academy to L.B., and a grant from the Deutsche Forschungsgemeinschaft (SPP 2330 project number 464976318) to C.S.

## Supplemental Information

### SUPPLEMENTARY TEXT

#### Growth and infection

*Haloferax gibbonsii* LR2-5 was grown aerobically in liquid modified growth medium (MGM) at 37 °C as described by Schwarzer et al. (2023). This medium contains 18% artificial salt water (SW) and yeast extract (0.1% w/v Oxoid) and peptone (0.5% w/v Oxoid) (Nuttall & Dyall-Smith, 1993). *H. gibbonsii* LR2-5 was grown in liquid medium until OD ∼1. Then the culture was diluted and divided over 6 ×100 mL Erlenmeyers with MGM medium to a theoretical OD of 0,05. After 24 hours of growth, when the OD was ∼0.5, to 3 of the 6 Erlenmeyer, HFTV1 particles were added to a multiplicity of infection (M.O.I.) of 10. HFTV1 stocks were prepared as described in Schwarzer et al 2023 (Schwarzer et al., 2023). The cultures were kept at 37 °C and incubated aerobically. Samples for RNA extraction were taken at 5, 20, 60, 120, 180 and 300 minutes post infection (p.i.) from infected and non-infected cultures in three biological replicates. At each time point, 2 mL samples were collected and immediately centrifuged at 3000 × g for 2 min. The supernatant was discarded and the cell pellet was flash frozen in liquid nitrogen and stored at −80°C.

#### Probe design, synthesis and chemical labeling

A total of 57 polynucleotide probes targeting the 38 kb viral genome of HFTV1 (NCBI accession no. NC_062739.1) were designed using the genePROBER Software (gene-prober.icbm.de/) (Moraru, 2021). Sequences of the 300-bp polynucleotides are listed in Supplemental Table 1. Selected probes were chemically synthesized as gBlocks® gene fragments (500 ng per polynucleotide) by IDT (Integrated DNA Technologies, San Jose, CA, USA), resuspended in 5 mM Tris, 1 mM EDTA, pH 8.0 prior to labeling with the ULYSIS™ Alexa Fluor™ 594 Nucleic Acid Labelling Kit (Thermo Fisher Scientific, Waltham, MA, USA) according to the manufacturer’s instructions. 2 µg HFTV1 targeting polynucleotides were labeled in a single reaction and subsequently purified using Micro Bio-Spin® Columns with Bio-Gel® P-30 (Bio-Rad Laboratories Inc., Hercules, CA, USA). Labeling efficiency was determined by spectrophotometric measurement using a NanoPhotometer® N50 (Implen, Munich, Germany) and calculated 9.12%. Probe specificity was checked against uninfected cells to exclude the possibility of nonspecific binding to cell structures.

#### Virus targeting direct-geneFISH (virusFISH)

Fluorescence in situ hybridization (FISH) was performed in accordance with the direct-geneFISH protocol of Barrero-Canosa and Moraru (2021) . Virus infections were performed as described in growth and infection using an M.O.I.. of 10. Cells were fixed in a paraformaldehyde solution (PFA) (Thermo Fisher GmbH, Kandel, Germany) as described in the “Core” direct-geneFISH protocol. 10 µl of fixed cells were spotted within a Silicone IsolatorTM (Grace Bio-labs, Bend, OR, USA) on a SuperfrostTM plus adhesion microscopy slides (Epredia Netherlands B.V, Essendonk, The Netherlands), air-dried and dehydrated by washing in ethanol with 50, 70 and 100% (v/v) for 10 minutes at room temperature. Haloarchaea have an S-layer consisting mainly of glycoproteins and no peptidoglycan, which makes them naturally more permeable and does not require an additional step to permeabilize. The samples were overlayed with a solution of 80 µl hybridization buffer containing 35 % (v/v) Formamide, 5 × SSC buffer (saline sodium citrate, pH 7.0), 20% (w/v) dextran sulfate, 20 mM EDTA, 0.25 mg mL^−1^ sheared salmon sperm DNA, 0.25 mg mL^−1^ yeast RNA, 1 × blocking reagent, 0.1% (v/v) sodium dodecyl sulfate, nuclease-free water, and probes were added at a final concentration of 30 pg µl^-1^ for each polynucleotide probe. To denature probes and templates, the samples were incubated at 92 °C for 40 min, followed by 2 h hybridization in a humidity chamber at 46 °C. Following hybridization, samples were rinsed in wash buffer I (2x SSC, 0.1% SDS) for 5 minutes at 48 °C, and then in wash buffer II (0.1x SSC, 0.1% SDS) for 30 minutes at 48°C. Finally, the samples were washed for 25 min in 1 × PBS and 1 min in ultra-pure water and air-dried. Cells were counterstained for 10 minutes in the dark with 10 µl of a mixture of SlowFade™ Diamond Antifade Mountant with 4′,6-diamidin-2-phenylindole (DAPI) (Thermo Fisher Scientific, Waltham, MA, USA) and covered with a #1.5 high-precision coverslip (Paul Marienfeld GmbH, Lauda-Königshofen, Germany). Images were acquired with a 100 × Plan-Apochromat oil objective (Numerical Aperture, NA = 1.46) on a Zeiss Axio Observer 7 inverted fluorescence microscope (Carl Zeiss Microscopy GmbH, Jena, Germany) equipped with a Colibri 7 LED illumination system and a Prime BSI Express sCMOS camera (Teledyne Photometrics, Tucson, AZ, USA). Phase contrast was imaged with an exposure time of 20 ms. DAPI was excited using an LED at 385 nm, and emission was collected using a BP 460/50 nm filter with an exposure time of 20 or 70 ms. Alexa Fluor 594 was excited using an LED at 555 nm, and fluorescence was detected using a BP 630/75 nm emission filter with an exposure time of 20 ms. Image acquisition and processing were performed using Zeiss ZEN (Blue Edition) software (version 3.5).

#### Bioinformatic genome analysis

Pharokka v 1.7.5 (Bouras et al., 2022) was used to update the annotation of the HFTV1 reference genome (NC_062739.1). More specifically PHANOTATE (McNair et al., 2019) was used to predict coding sequences (CDS), while tRNAscan-SE 2.0 (Chan et al., 2021) was used to predict tRNAs. The respective functional annotations were assigned by matching each CDS to the PHROGs (Terzian et al., 2021) VFDB (Chen et al., 2005) and CARD (Alcock et al., 2019) databases using MMseqs2 (Steinegger & Söding, 2017) and PyHMMER (https://github.com/althonos/pyrodigal-gv). By using Mash (Ondov et al., 2016) contigs were matched to their closest hit in the INPHARED database (Cook et al., 2021). The functional annotation was refined using Phold (https://github.com/gbouras13/phold) which uses ProstT5 (Heinzinger et al., 2024) for the generation of protein structures for each CDS. Followed by a comparison against a database of predicted viral protein structures using FoldSeek (van Kempen et al., 2024). The resulting output plots were created using pyCirclize (https://github.com/moshi4/pyCirclize).

#### Expression analysis

The RNA sequencing, including the rRNA depletion and cDNA synthesis, was performed by Vertis Biotechnologie AG (Freising, Germany) and the pooled cDNA was sequenced on an Illumina 500 system using 1×75 bp read length. On average 10.5 million reads per sample were obtained. The FastQ files were run through a BowTie2-SamTools pipeline and the resulting BAM files were analysed using SeqMonk V1.48.1 in which the *H. gibbonsii* LR2-5 strain (Genome assembly ASM1496974v1) or Haloferax tailed virus 1 (HFTV1, Genome assembly ASM420877v1) were used as reference genomes. All *H. gibbonsii* LR2-5 genes were analysed, using SeqMonk, in an intensity difference statistical test (DESeq2) in which a statistical difference of below 0.05 was used (p<0.05). The EggNOGv5.0 (http://eggnog-mapper.embl.de) was subsequently used to functionally classify all *H. gibbonsii* LR2-5 genes in the various archaeal clusters of orthologous genes (arCOGs).

#### Swimming analysis with time-lapse microscopy

For analysis of swimming behaviour, an overnight culture was diluted in MGM medium, to a theoretical OD of 0,05. After 24 hours, when the OD was ∼0,5, the culture was infected with HFTV1 to a M.O.I. of 10. Samples were taken at different time points post infection. In all, 1 mL of each culture was placed in a round DF 0.17 216 mm microscopy dish (Bioptechs, Butler, PA, USA) and observed with an 40 × Plan-Apochromat objective (Numerical Aperture, NA = 0.95) in the PH3 mode with a Zeiss Axio Observer 7 (Carl Zeiss Microscopy GmbH, Jena, Germany) equipped with a heated XL Incubator combined with the S1 temperature control, heated to 45 °C running Zen 3.5 software.

### SUPPLEMENTARY DATA

#### Supplemental Tables

**Supplemental Table S1:**
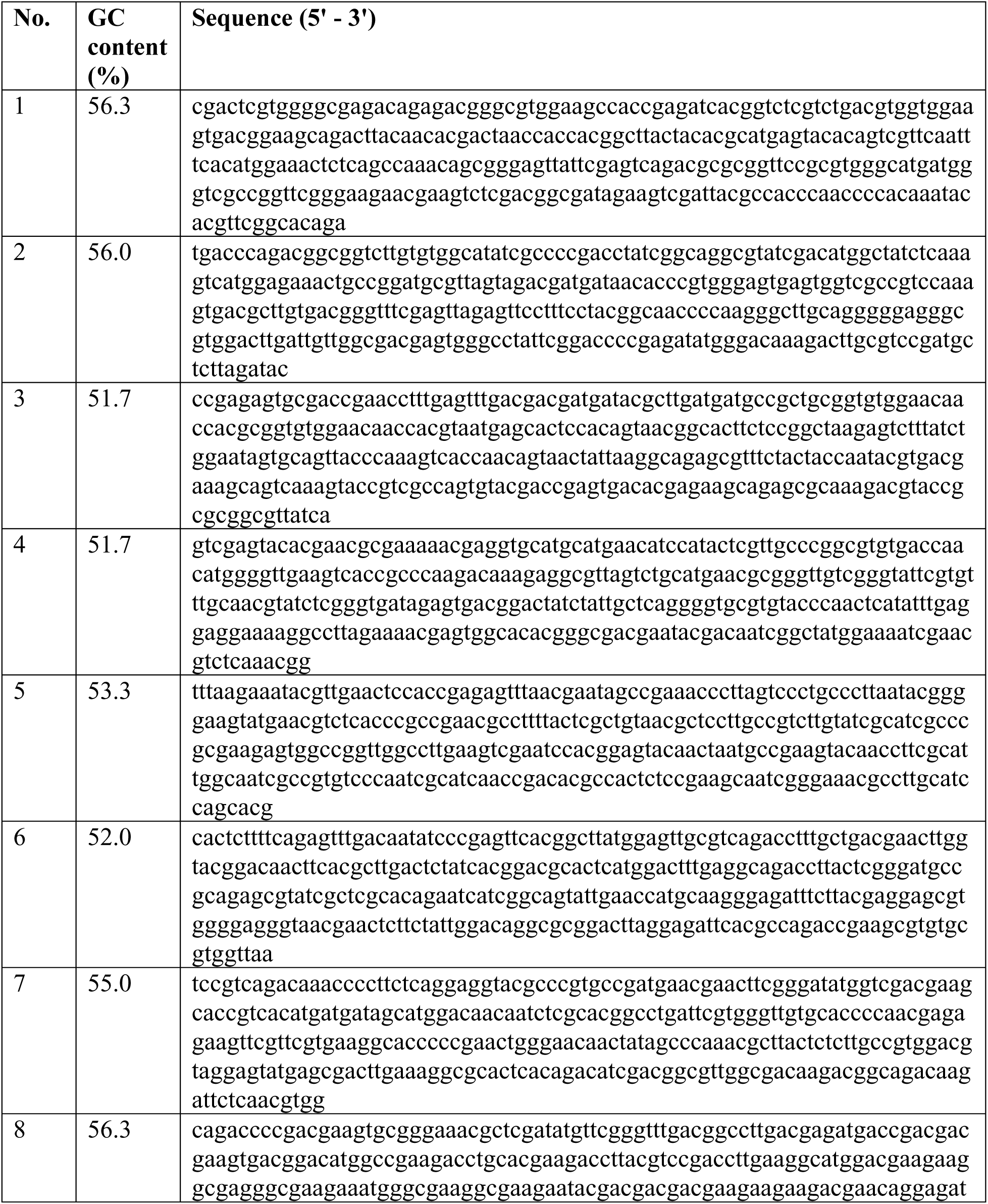
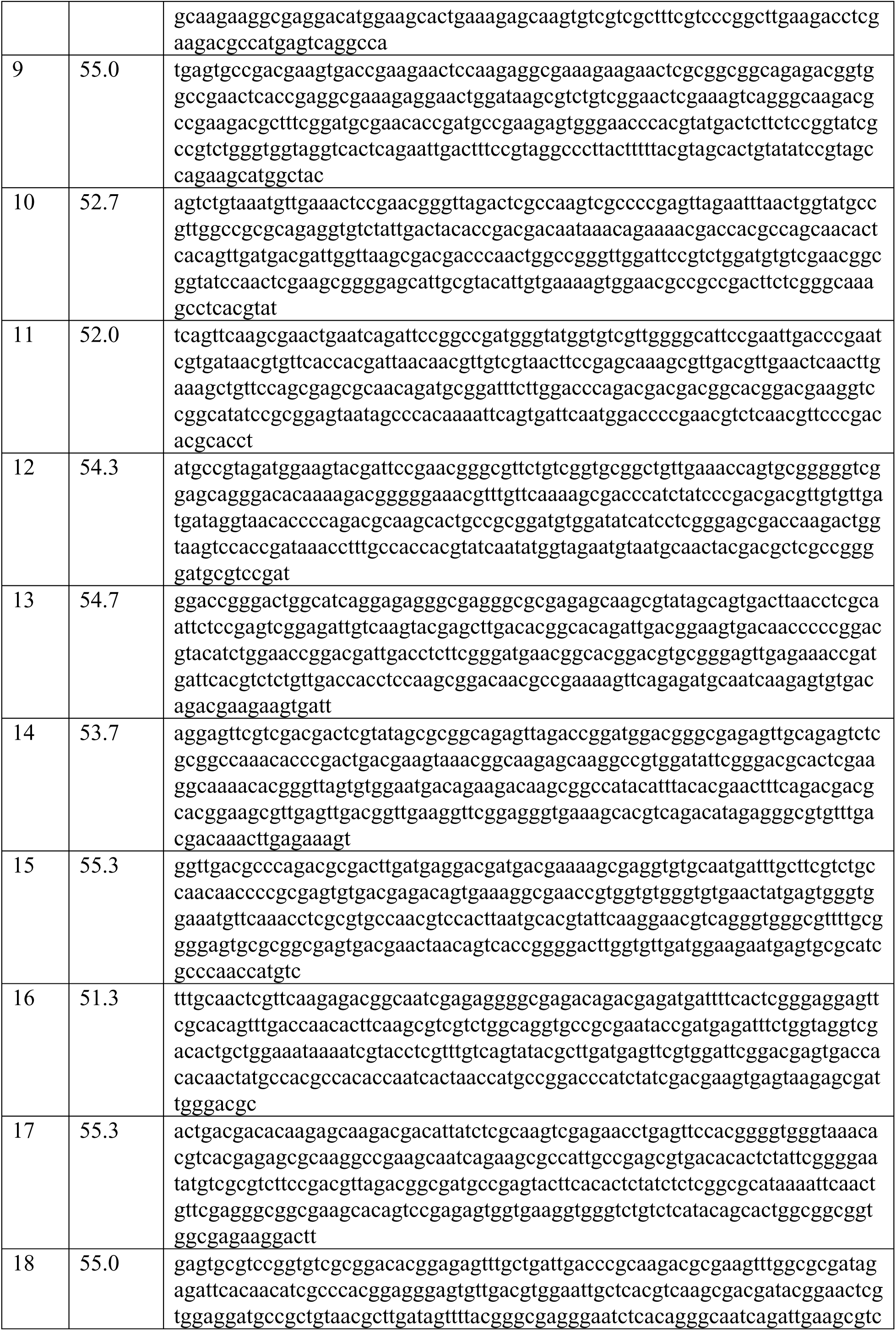
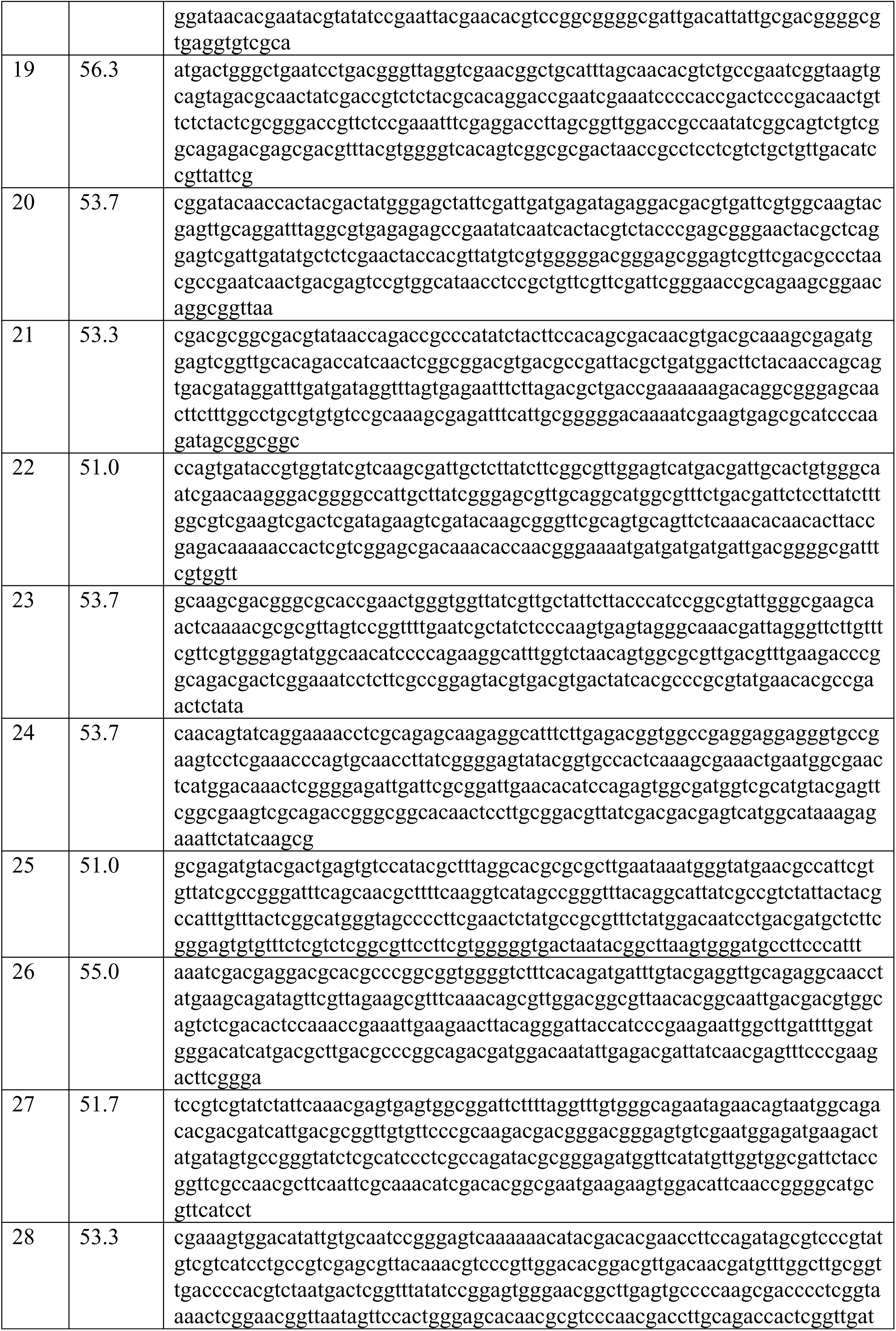
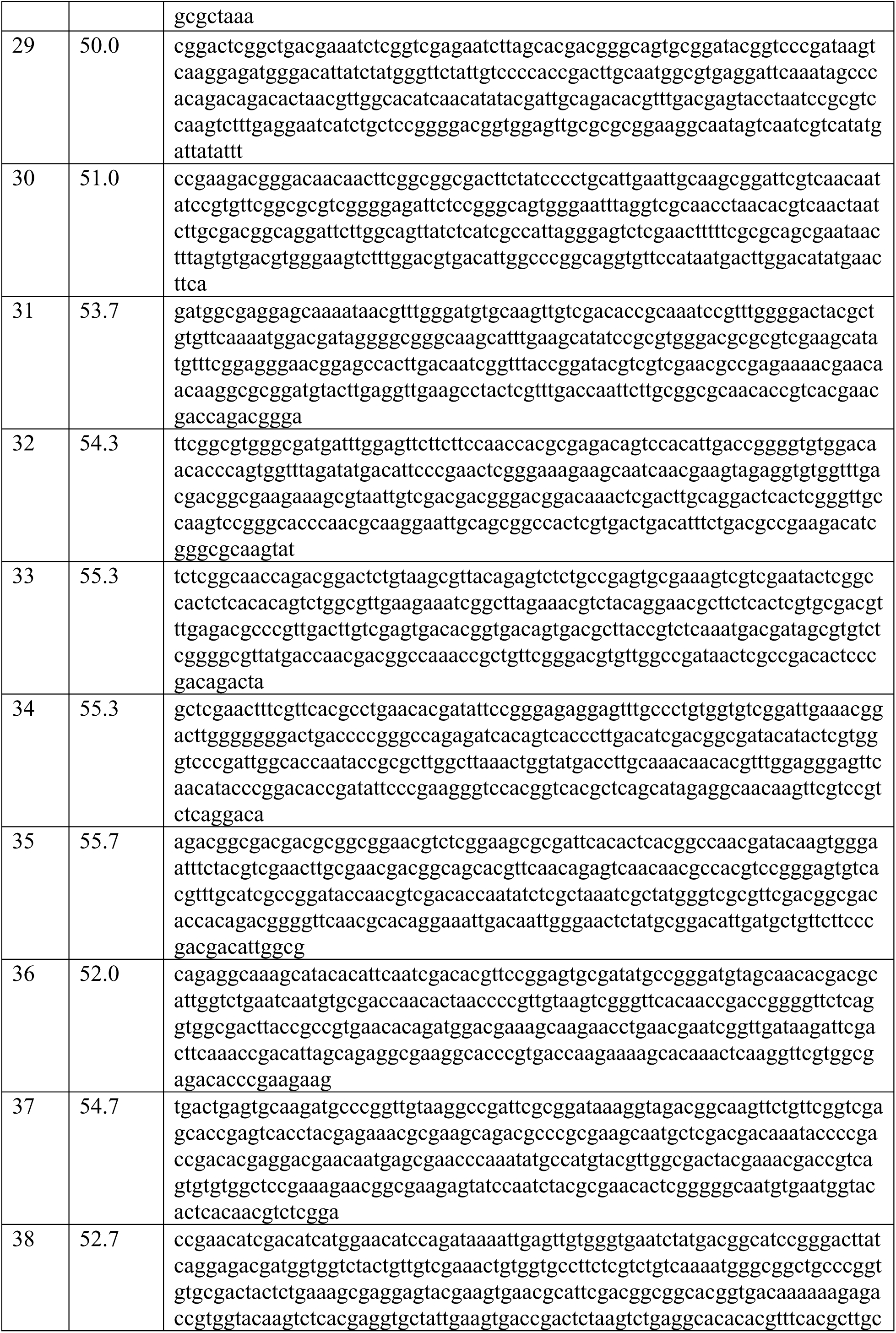
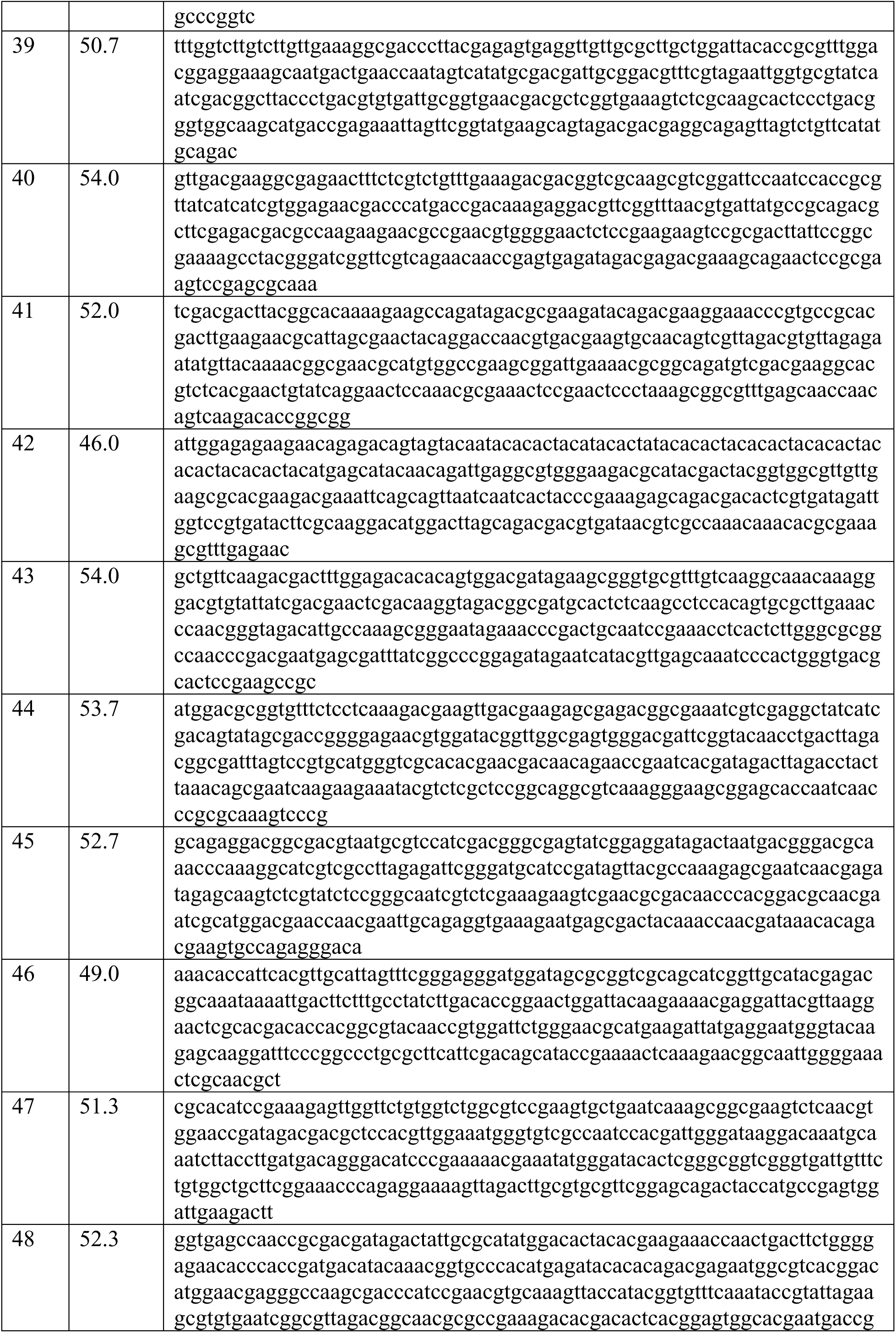
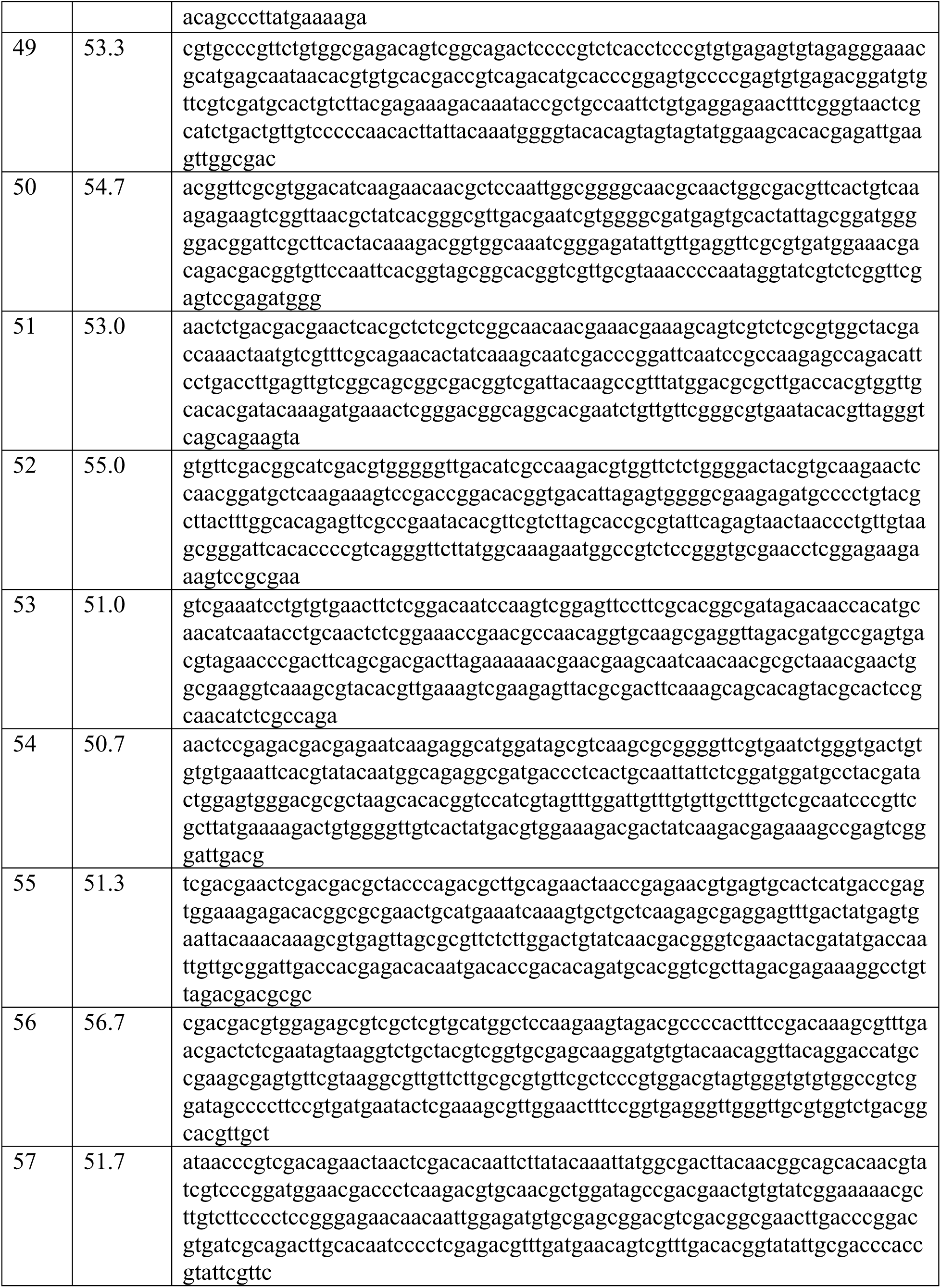
Polynucleotides used for virus targeting direct-geneFISH with HFTV1.

**Supplemental Table S2:** Coverage values for sense and antisense transcripts across the HFTV1 genome, based on RNA-seq data.

| Sense transcript |  |  |  |  |  |  |
| --- | --- | --- | --- | --- | --- | --- |
| Coverages | T5 | T20 | T60 | T120 | T180 | T300 |
| TSS gp01 | 1 | 12 | 54 | 100 | 97 | 64 |
| TSS post gp02 | 0 | 3 | 11 | 100 | 92 | 58 |
| TSS gp06 | 1 | 11 | 100 | 75 | 57 | 49 |
| TSS gp06a | 9 | 67 | 100 | 40 | 22 | 17 |
| TSS gp13 | 0 | 1 | 3 | 100 | 76 | 45 |
| TSS gp14 | 0 | 1 | 7 | 100 | 92 | 98 |
| TSS non coding RNA | 0 | 4 | 21 | 59 | 100 | 54 |
| TSS gp15 | 0 | 1 | 8 | 68 | 95 | 100 |
| TSS gp18 | 0 | 0 | 2 | 82 | 100 | 57 |
| TSS gp20 | 0 | 1 | 3 | 100 | 96 | 77 |
| TSS gp21 | 1 | 12 | 33 | 100 | 61 | 26 |
| TSS gp25 | 0 | 0 | 0 | 19 | 100 | 96 |
| TSS gp26 | 1 | 2 | 15 | 96 | 93 | 100 |
| TSS gp29 | 1 | 5 | 17 | 100 | 86 | 75 |
| TSS gp30 | 0 | 1 | 7 | 100 | 100 | 62 |
| TSS gp33 | 0 | 1 | 1 | 29 | 96 | 100 |
| TSS gp35 | 0 | 1 | 8 | 100 | 65 | 41 |
| TSS gp38 | 1 | 4 | 22 | 100 | 90 | 60 |
| TSS gp39 (anti gp38) | 0 | 2 | 17 | 87 | 99 | 100 |
| TSS gp40 | 1 | 4 | 8 | 71 | 75 | 100 |
| TSS gp42 | 0 | 2 | 11 | 100 | 60 | 40 |
| TSS gp45 | 1 | 4 | 72 | 100 | 52 | 37 |
| TSS gp46a | 3 | 27 | 100 | 14 | 4 | 3 |
| TSS gp46b | 1 | 13 | 100 | 22 | 12 | 17 |
| TSS gp46 | 4 | 32 | 100 | 11 | 12 | 20 |
| TSS gp51 | 0 | 0 | 18 | 60 | 94 | 100 |
| TSS gp52 | 4 | 60 | 100 | 24 | 21 | 19 |
| TSS post gp56 | 1 | 1 | 15 | 67 | 100 | 75 |
| TSS pre gp61 | 0 | 3 | 18 | 83 | 100 | 75 |
| TSS gp62 | 12 | 100 | 46 | 29 | 31 | 23 |
| TSS gp63 | 17 | 100 | 41 | 20 | 10 | 9 |
| TSS gp64 | 11 | 100 | 45 | 39 | 28 | 18 |
| TSS pre gp66 | 0 | 4 | 27 | 100 | 90 | 55 |
| TSS post gp66 | 0 | 0 | 11 | 100 | 93 | 74 |
| TSS gp68 | 0 | 3 | 68 | 100 | 48 | 20 |
| **Antisense transcript** |  |  |  |  |  |  |
| **Coverages** | **T5** | **T20** | **T60** | **T120** | **T180** | **T300** |
| TSS anti gp6a | 0 | 0 | 7 | 60 | 100 | 75 |
| TSS anti gp13 | 0 | 2 | 8 | 40 | 100 | 76 |
| TSS anti gp13 | 0 | 1 | 4 | 44 | 100 | 75 |
| TSS anti gp18 | 1 | 8 | 25 | 86 | 100 | 90 |
| TSS anti gp20 | 1 | 4 | 9 | 58 | 100 | 76 |
| TSS anti gp20 | 0 | 0 | 0 | 4 | 33 | 100 |
| TSS anti gp24 | 0 | 2 | 18 | 55 | 79 | 100 |
| TSS anti gp25 | 2 | 2 | 12 | 71 | 97 | 100 |
| TSS anti gp26 | 2 | 11 | 17 | 47 | 86 | 100 |
| TSS anti gp33 | 0 | 1 | 5 | 39 | 100 | 95 |
| TSS anti gp41 | 1 | 37 | 24 | 100 | 87 | 49 |
| TSS anti gp42 | 0 | 13 | 21 | 82 | 86 | 100 |
| TSS anti gp44 | 0 | 1 | 9 | 67 | 96 | 100 |
| TSS anti gp47 | 1 | 1 | 7 | 51 | 97 | 100 |
| TSS anti gp48 | 1 | 3 | 9 | 100 | 96 | 69 |
| TSS anti gp50 | 0 | 3 | 6 | 59 | 100 | 100 |
| TSS anti gp51 | 0 | 0 | 2 | 15 | 80 | 100 |
| TSS anti gp56 | 0 | 0 | 3 | 59 | 100 | 56 |
| TSS anti gp 58 | 1 | 2 | 13 | 90 | 100 | 62 |
| TSS anti gp59 | 0 | 2 | 9 | 61 | 100 | 85 |
| TSS anti gp60 | 1 | 3 | 22 | 100 | 88 | 52 |
| TSS anti gp65 | 1 | 5 | 8 | 66 | 100 | 58 |

#### Supplementary Figures

**Supplemental Figure 1.**
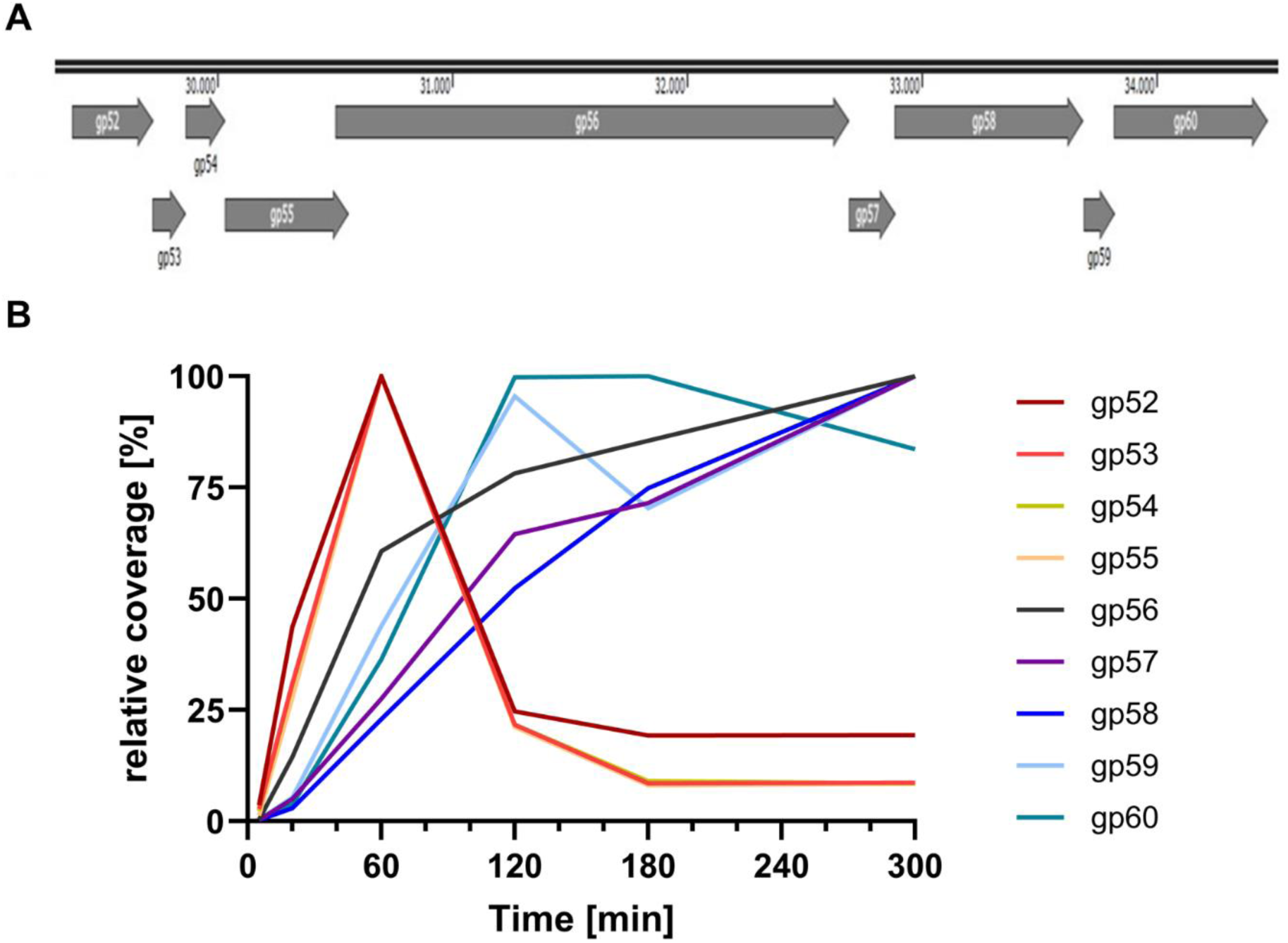
Gene cluster structure (A) and percentage of total read counts (B), of the HFTV1 viral gene cluster and *gp52* - *gp60*. Depicted in black/grey *gp52* – *gp55* and in green *gp56* - *gp60*.

**Supplemental Figure 2:**
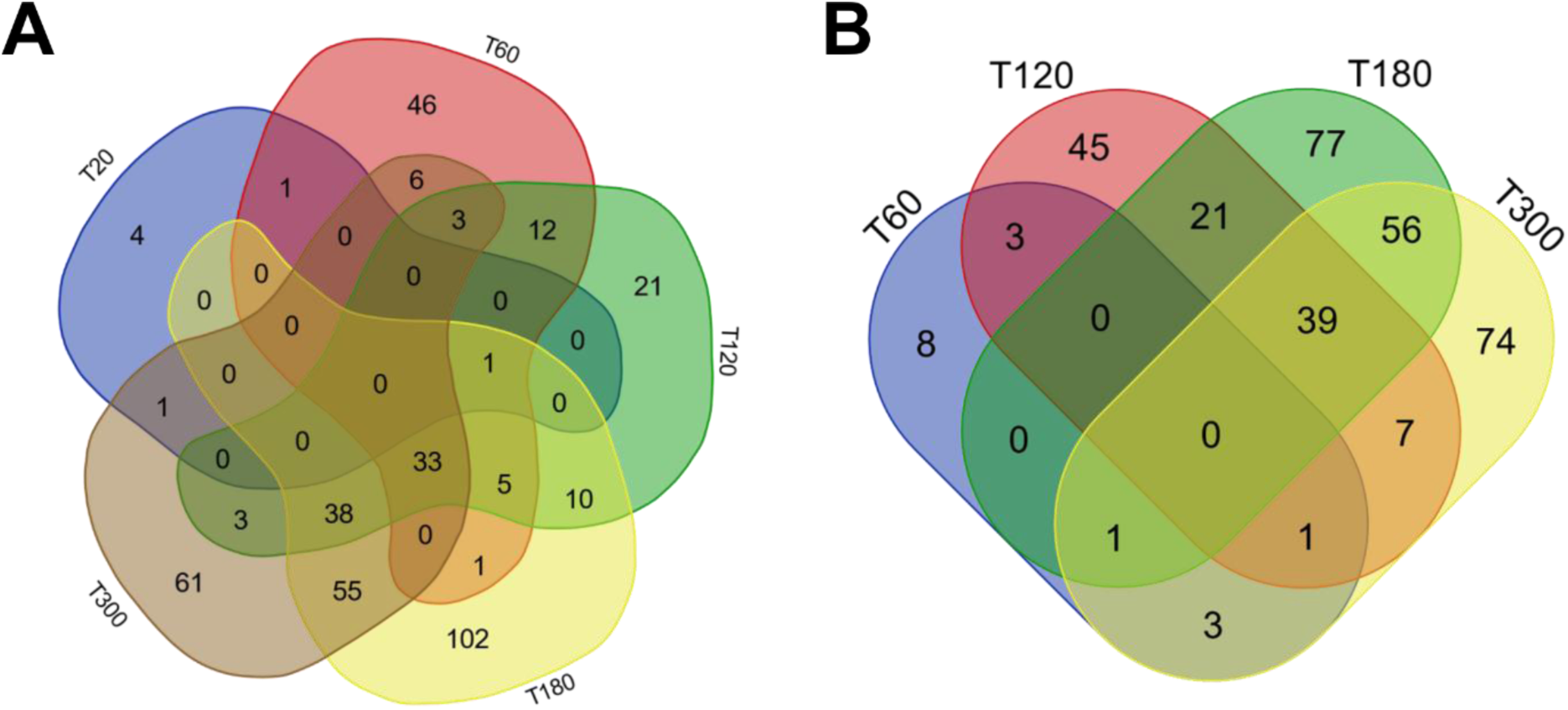
Venn diagrams of differentially expressed genes in *H. gibbonsii* in response to HFTV1 infection. (A) Upregulated genes and (B) Downregulated genes at 5, 20, 60, 120, and 300 minutes post-infection (p.i.). Each diagram represents the number of differentially expressed genes (DEGs) at each time point and their overlap, highlighting shared and unique transcriptional responses across the infection timeline.

### SUPPLEMENTARY VIDEO

**Supplemental Movie 1**. Time-lapse light microscopy of H. gibbonsii cells under standard conditions. Time-lapse phase-contrast microscopy of Haloferax gibbonsii cells under untreated conditions. Imaging was conducted at 45 °C, over a duration of 1 minute at 0.8 frames per second (fps).

**Supplemental Movie 2**. Time-lapse light microscopy of H. gibbonsii post HFTV1 infection. Time-lapse phase-contrast microscopy of H. gibbonsii 5 hours after infection with HFTV1 at a multiplicity of infection (M.O.I.) of 10. Imaging was performed at 45 °C, over a duration of 1 minute at 0.8 frames per second (fps).

